# Loss of Gαq reshapes key fibroblast traits and drives matrix remodeling and aggressive progression of oral cancer tumors

**DOI:** 10.1101/2025.03.12.642811

**Authors:** Inmaculada Navarro-Lérida, Raquel Huertas-Lárez, Gemma Paredes-García, Dafne García-Mateos, María Isabel Jiménez-López, Juan Antonio López, Miguel A. del Pozo, Saúl Álvarez-Teijeiro, Llara Prieto-Fernández, Juana María García-Pedrero, Federico Mayor, Catalina Ribas

**Affiliations:** Departamento de Biología Molecular, Instituto Universitario de Biología Molecular IUBM-Universidad Autónoma de Madrid and Centro de Biología Molecular Severo Ochoa (Universidad Autónoma de Madrid-CSIC).28049 Madrid. Spain; Instituto de Investigación Sanitaria Hospital Universitario La Princesa, 28006 Madrid. Spain; Centro de Investigación Biomédica en Red de Enfermedades Cardiovasculares (CIBERCV). Madrid. Spain; Conexión Cáncer-CSIC; Centro Andaluz de Biología Molecular y Medicina Regenerativa. Sevilla. Spain; Proteomics Unit; Centro Nacional de Investigaciones Cardiovasculares (CNIC). Madrid. Spain; Mechanoadaptation and Caveolae Biology Laboratory, Cell and Developmental Biology Area, Centro Nacional de Investigaciones Cardiovasculares (CNIC). 28029, Madrid. Spain; Instituto de Investigación Sanitaria del Principado de Asturias (ISPA), 33011 Oviedo, Spain; Instituto Universitario de Oncología del Principado de Asturias, Universidad de Oviedo, 33006 Oviedo, Asturias, Spain; Spanish Biomedical Research Network in Cancer (CIBERONC), Instituto de Salud Carlos III, 28029 Madrid, Spain

**Keywords:** Tumor stroma, CAFs, Head and Neck Squamous Cell Carcinoma (HNSCC), Gαq protein, exosomes

## Abstract

Head and neck squamous cell carcinoma (HNSCC) is a highly aggressive cancer, with limited therapeutic options and a high mortality rate, primarily due to metastasis and recurrence. Tumor-stroma interactions, and namely cancer-associated fibroblasts (CAFs), are pivotal in shaping HNSCC progression. CAFs remodel the extracellular matrix (ECM) and secrete factors and vesicles that promote tumor growth and metastasis. The interplay between autophagy and endosomal/exosomal pathways has been suggested to regulate cellular secretory functions, but their potential involvement in HNSCC progression remains poorly understood. Since we have recently uncovered Gαq as a key modulator of autophagy, we have investigated the impact of Gαq loss on fibroblast functionality and on its crosstalk with oral HNSCC cells.

We report that the absence of Gαq rewires murine embryonic fibroblasts towards CAF-like traits, leading to an increased pro-tumorigenic capacity of co-cultured human oral cancer cells through enhanced collagen I deposition and ECM remodeling. Strikingly, fibroblasts lacking Gαq display a shift in the balance of intracellular trafficking, degradative and secretory pathways. Exosomes released from Gαq-deficient fibroblasts show a marked enrichment in tumor-growth factor receptors and can facilitate aberrant tumor growth of HNSCC cells. Gαq-silenced fibroblasts promote the formation of “railroad-tracks” structures around HNSCC cells, enhancing their migratory and invasive capabilities both *in vitro* and *in vivo*, and reduced Gαq expression in human HNSCC CAFs correlates with enhanced tumor progression. Overall, our data put forward Gαq as a key regulator of the HNSCC tumor microenvironment by modulating fibroblast plasticity and functionality.

## Introduction

Tumors are constantly seeking alternative adaptive mechanisms to ensure progression, and it is now recognized that the changing tumor microenvironment (TME) and the mechanisms of bidirectional communication between cancer and stromal cells are critical in this quest. Consisting of a complex meshwork of extracellular matrix (ECM) proteins, secreted factors and different types of cells, the TME is a dynamic entity that provides both biophysical and biochemical cues that influence multiple parameters of cancer growth and aggressiveness ^1^. Key regulators of the TME are cancer-associated fibroblasts (CAFs), a heterogeneous and plastic population of fibroblasts that are activated in pathological contexts and drive tumor progression through various mechanisms ^2,3^.

Although most solid tumors are highly dependent on CAFs, this cell population is particularly relevant in head and neck squamous cell carcinoma (HNSCC), a severe and complex malignancy ^4^. High density of biologically heterogenous CAFs has been associated with poor prognostic features (proliferation, migration and invasion) and higher rates of local recurrence in HNSCC ^5^. Remarkable elevated amounts of CAFs, up to 80-90% of tumor volume, occur in later stage HNSCC associated with a higher metastatic phenotype ^6,7^

From a molecular perspective, CAFs can express high levels of several specific markers, such as the platelet-derived growth factor receptor α/β (PDGFR α/β) ^8–11^. Interestingly, dysregulation of spatio-temporally controlled PDGFR-induced signaling has emerged as a critical mechanism governing tumor growth and survival. Within the context of HNSCC, the overexpression of PDGF and its receptor has been associated with oral tumorigenesis and poor prognosis ^12,13^. However, the underlying mechanisms regulating PDGFR modulation and dynamics remain poorly understood.

CAFs play a critical role in shaping the mechanical tumor microenvironment by depositing and remodeling ECM components ^14,15^. Besides, CAFs secretome is emerging as a key regulator of tumor progression, by enabling communication with cancer cells and other cells in the TME via both soluble factors and exosomes. As a membranous nanovesicle system, exosomes can mediate the intercellular transport of multiple molecules such as proteins, nucleic acids, and lipids, which can influence a variety of cell functions both locally and remotely ^16,17^. Membrane proteins reported to be present on specific exosomes include the epidermal growth factor receptor (EGFR), glycosylphosphatidylinositol (GPI), HER2, platelet-derived growth factor receptor (PDGFR), among many others ^17,18^. These receptors may directly activate pro-survival signaling pathways in the recipient cells, thus promoting resistance to chemotherapy. ^19,20^

Of note, emerging evidence shows a direct correlation between exosomes and the aggressiveness of HNSCC ^21,22^ and highlights the potential role of the interplay between autophagy, a self-catabolic process, and exosome biogenesis and secretion as a pivotal mechanism in regulating this type of tumor. Both cellular processes depend on the regulation of the endolysosomal system and rely on the convergence of degradative and secretory regulators ^23,24^. However, the mechanisms underlying the specific sorting of certain growth factor receptors to exosomes and the crosstalk between secretory and autophagy pathways in pathological contexts remain unclear.

G-protein-coupled receptors (GPCR) play a fundamental role in the control of cellular homeostasis. However, aberrant expression, function, or mutations of GPCRs and/or their intracellular signaling networks also significantly contribute to many facets of tumorigenesis, often acting as driving oncogenes themselves ^25,26^. Interestingly, recent studies suggest the potential involvement of GPCRs and their downstream signaling pathways in the biogenesis, secretion, homing, and uptake of extracellular vesicles ^27^. Within this system, mutated versions of the Gαq protein are present in different tumor types, particularly in uveal and cutaneous melanomas ^28,29^. Gαq signaling drives tumorigenesis by mechanisms related to the activation of its canonical effectors, such as phospholipase C-beta, as well as via an emerging complex interactome involving a novel effector region capable of interacting with proteins with PB1 domains ^30^. Remarkably, we have reported that Gαq is a crucial regulator of the autophagy process in mouse fibroblasts, by mechanisms involving the modulation of active mTORC1 complexes via this novel non-canonical effector region ^31^. Therefore, we set to explore whether Gαq may play a more general role in the crosstalk between endolysosomal and secretory pathways and its potential impact on the tumor stroma in HNSCC pathological contexts.

Here, we demonstrate that the absence of Gαq in fibroblasts leads to their activation, resulting in matrix remodeling and contractile features comparable to malignant CAFs, and to marked alterations in protein trafficking and degradation, which results in aberrant cargo sorting to released exosomes of several tumor growth factor receptors, such as Platelet-derived-growth-factor receptor (PDGFR), a key driver of HNSCC progression. Consistently, Gαq-KO fibroblasts or their derived exosomes foster oral cancer cell growth and invasiveness. Our data put forward stromal Gαq as a new player in the modulation of the HNSCC tumor microenvironment by controlling fibroblast plasticity and functionality as integrator of ECM remodeling and exosome/autophagic flux control.

## Results

### Gαq deficiency in fibroblasts promotes Cancer-Associated Fibroblasts (CAFs) traits

To address the potential role of Gαq at the stromal level, we assessed the impact of Gαq absence on the activation status of Mouse Embryonic Fibroblasts (MEFs). After 24 hours of seeding, we compared the expression of key CAF-related biomarkers between wild-type (WT) and Gαq knockout (GαqKO) MEFs. The lack of Gαq expression, as confirmed by both Western blot (Fig.1A) and immunofluorescence (Fig.S1A-B), was associated with a significant upregulation of most of the CAFs markers assessed (PDGFR, Caveolin1, PTRF (also referred to as Cavin-1), FSP1 and osteopontin), except for α-Smooth-Muscle-Actin (α-SMA).

**Figure 1.**
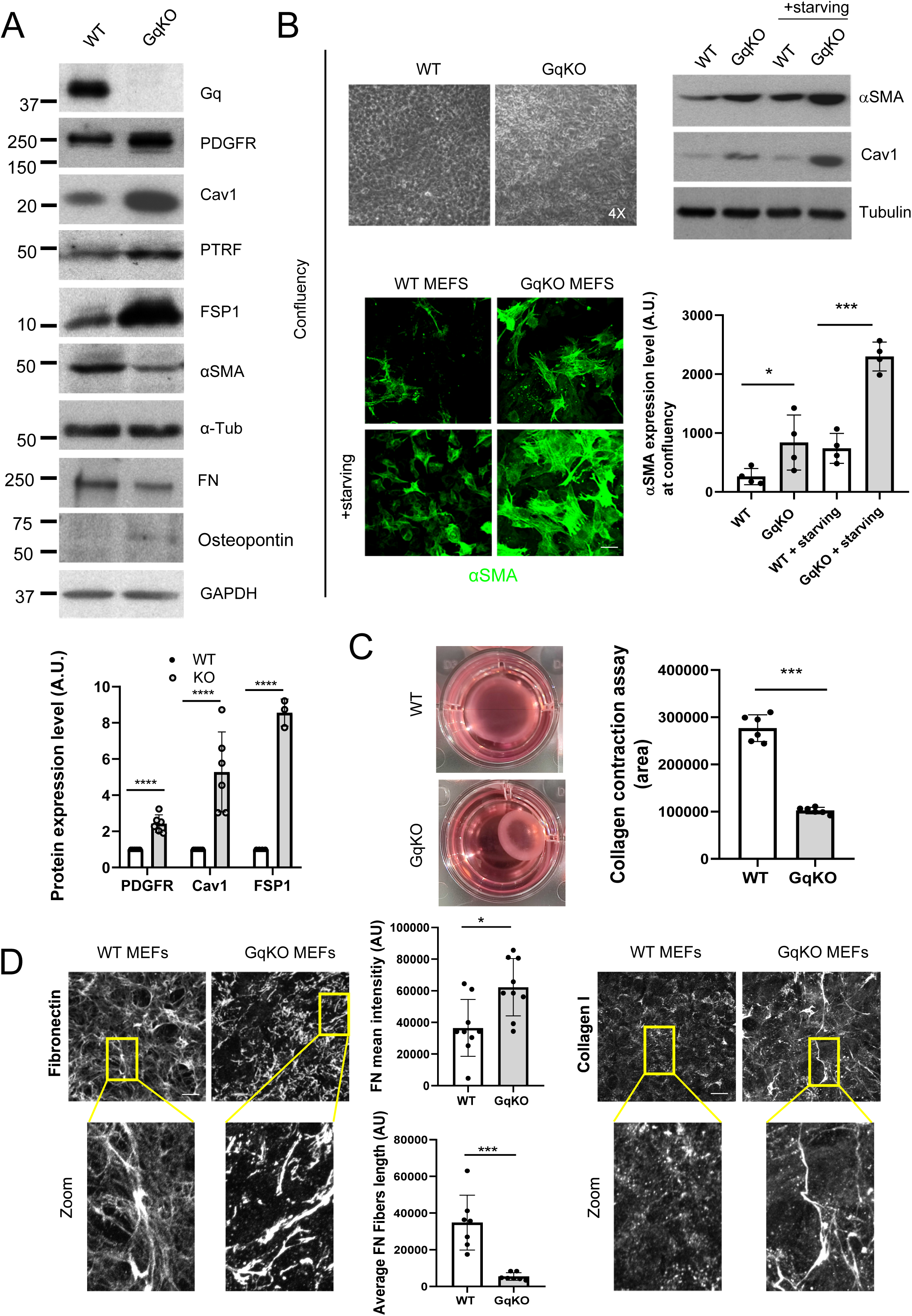
Lack of Gαq in fibroblasts results in Cancer-Associated Fibroblasts (CAFs) features. (a) Western blot assessing fibroblast activation markers in wild-type and GαqKO MEFs after 24 h in culture. Tubulin and GAPDH were used as loading controls. A comparison of the protein expression levels of the markers PDGFR, Caveolin-1, αSMA, and FSP1 is shown in the graph below (mean±SD of 3-6 experiments). (b) αSMA protein expression at high confluency in wild-type and GαqKO MEFs under baseline or starvation conditions (mean±SD, n≥3). Confocal images of αSMA taken under the same circumstances as in b are displayed below. The chart shows the amounts αSMA at confluent conditions (mean±SD, n=4). (c) Collagen gel contraction assay. Representative photomicrographs show collagen gel changes of MEFs wild-type or lacking Gαq expression. Plot represents contraction indicated as percentage of the initial gel surface area (mean±SD, n=6). (d) Z stack projection of confocal microscopy images showing Fibronectin (left panels) and Collagen I (right panels) in matrix deposited by wild-type or GαqKO MEFs (scale bar, 10 µm). The charts represent average intensity (upper graph) and fiber length (lower graph) of FN in each case, (mean±SD, n=9). For all graphs, *, p < 0.05; **, p < 0.01; ***, p < 0.001.

Since molecular variations between CAFs with different levels of α-SMA have been reported ^32–34^ and differences in the expression of α-SMA seem to be critically influenced by different factors, including cell morphology and adhesion, we evaluated whether those features could underlie the observed slightly reduced levels of α-SMA in GαqKO MEFs after 24 hours in culture. Of note, assessment of cell morphology by phase contrast microscopy at this time point revealed that GαqKO fibroblasts displayed a slightly delayed cell attachment, a more apparent spindle-like shape (Fig. S1A), and a higher F-actin stress fiber pattern (Fig. S1B) compared to WT MEFs. To address the potential impact of differences in initial cell adhesion, we determined the expression levels of α-SMA at confluency, after a period of three to five days of culture (Fig. 1B). In the absence of apparent differences in cell morphology at this stage (Fig. 1B upper left panels), there was a significant shift to higher expression of the α-SMA marker in the GαqKO MEFs compared to WT cells under such confluent conditions, which was further enhanced upon starvation (Fig. 1B). Other key CAF markers like Cav1 and PDGFR showed an even higher overexpression compared to the 24 h culture period (Fig. S1C). Interestingly, under these confluent conditions, GαqKO MEFs displayed a much higher contractile capacity compared with WT cells, as indicated by collagen contraction assays (Fig.1C).

In addition to increased contractile properties, one of the main features of CAFs is their ability to produce large amounts of ECM proteins, such as collagens, proteoglycans, and glycoproteins ^35^. The ECM matrices deposited by WT versus GαqKO MEFs were thus analyzed to elucidate the functional implications of Gαq deficiency on this key characteristic. Fibroblasts lacking Gαq expression deposited a fibronectin (FN) fiber matrix different from that of WT cells, characterized by shorter and significantly thicker FN fibers as evidenced by fluorescent immunolabelling and confocal imaging (Fig. 1D, left images and graphs). Interestingly, this phenotype was partially mimicked in wild-type MEFs treated with YM254890, a specific Gαq inhibitor (Fig. S1D). Moreover, in the same experimental conditions, wild-type MEFs deposited almost no collagen I fibers, in contrast to the remarkable collagen I matrix noted in GαqKO MEFs (Fig. 1D, right images).

Overall, these data indicated that Gαq plays a relevant role in modulating fibroblast activation, since its absence triggers most features of CAFs.

### GαqKO fibroblasts drastically shape the architecture, proliferation, and invasion potential of oral tumor cells in both 2D and 3D co-culture models

To determine the potential influence of such pro-tumoral stromal features triggered upon Gαq loss, we established a contact co-culture system using fibroblasts (with or without Gαq) and human head and neck squamous carcinoma (HNSCC) cells. Thus, three different oral HNSCC cell lines (HN13, UMSCC47 and Cal27) were co-seeded with wild-type (WT), Gαq knockout (GαqKO) or Gαq knock-in (GαqKI, a lentiviral-reconstituted version of GαqKO MEFs expressing wild-type Gαq) MEFs. After 72 h of co-culture, 2D phase contrast images revealed a completely different pattern of tumor-fibroblast distribution depending on stromal Gαq expression. While the 3 different oral tumor cells remained “encapsulated” in the presence of either WT or GαqKI MEFs, the distribution detected in the presence of GαqKO MEFs was far more chaotic, with tumor cells and fibroblasts exhibiting an intertwined pattern with less clearly defined encapsulated structures (Fig. 2A). To better visualize such differential distribution and further explore the ECM deposition and matrix organization under these co-culture conditions, FN and collagen I staining was performed in co-cultures of Cal27 oral cancer cells with WT or GαqKO fibroblasts stably expressing GFP. Again, in the presence of WT MEFs we observed encapsulated oral tumor cells, which were evenly distributed and formed a fibronectin-rich capsule largely devoid of collagen fibers (Fig. 2B, upper panel). In contrast, GαqKO MEFs fostered an entirely distinct pattern in which tumor cells and fibroblasts intermingled (Fig. 2B, lower images in upper panel), leading to a partial reduction of tumor cell-cell contacts and higher cell scattering (Fig. S2A). The most striking feature of the matrix developed under these conditions was a robust distribution of collagen I fibers, which created well-defined “railroad-tracks” moving away from the tumor cells (Fig. 2B, lower panel).

**Figure 2.**
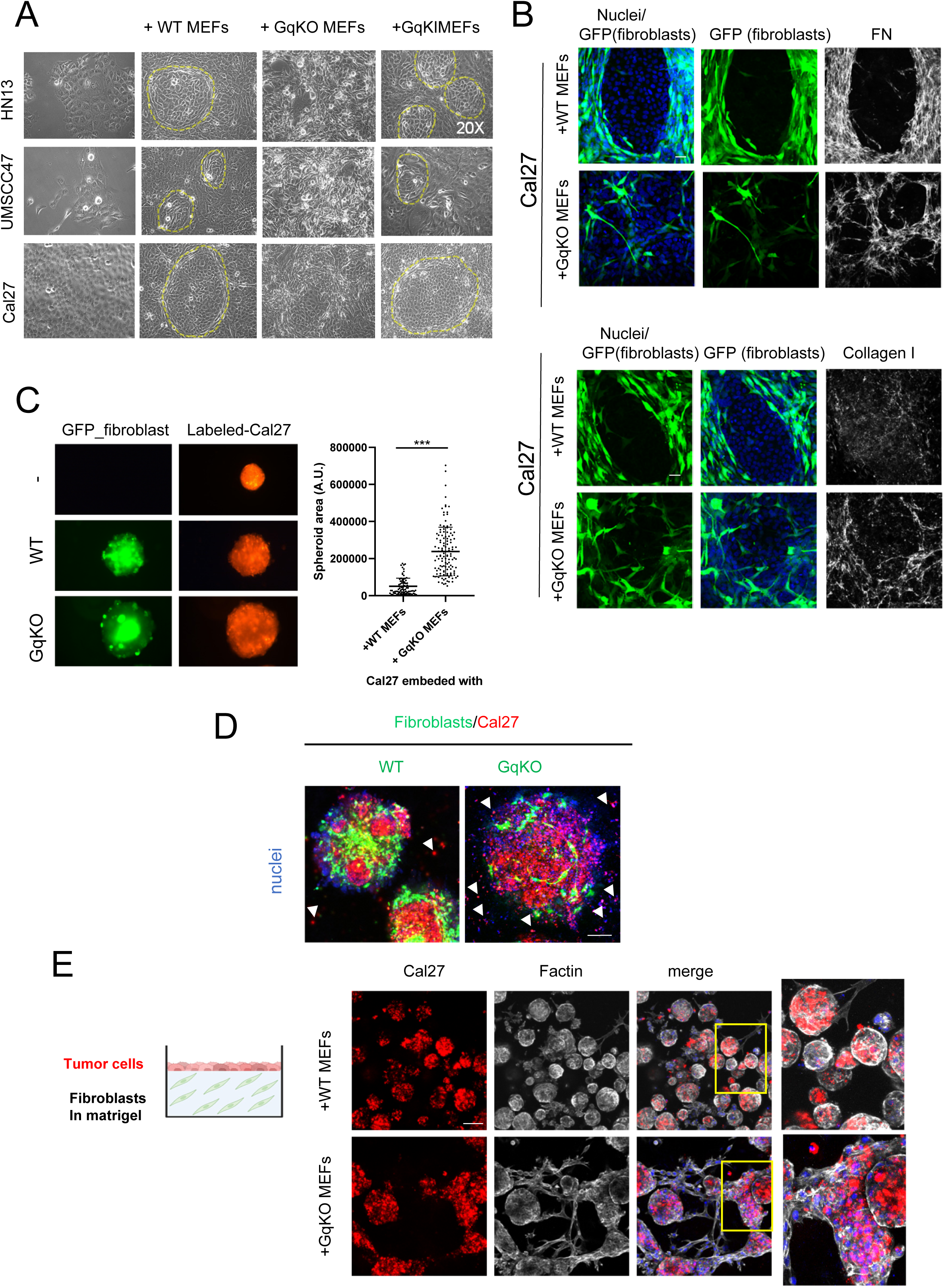
Impact of GαqKO Fibroblasts on oral Tumor Cell Organization, Growth, and Invasion in 2D and 3D co-culture Models. (a) Phase contrasts images (20X magnification) of 2D co-cultures of different human squamous cell carcinoma cell lines (Cal27, HN13 and UMSCC47) and either WT, GαqKO or GαqKI (lentiviral-reconstituted with the expression of wild-type Gαq) MEFs. (b) Analysis by confocal microscopy of the matrix deposited by GFP-labeled WT or GαqKOMEFs in co-culture with Cal27 cells. Fibroblasts are shown in green, FN or collagen I in grey and DAPI-labeled nuclei in blue. (c) Epifluorescence microscopy of tumor spheroid formation and growth formed in conditions of co-cultures of cell-tracker prelabeled Cal27 cells (red) with GFP-expressing WT or GαqKO MEFs (green). The plot represents the area of spheroid formation in each condition (mean± SD, n=82 spheroids per condition). (d) Invasion assay of Cal27 cells (red) spheroids generated in the presence of WT or GαqKO MEFs (green) embedded into Matrigel. Arrows point to invading cells (scale bar 150 µm). (e) The left scheme shows the assay protocol. Images show the Z-stack confocal analysis of an invasion assay of Cal27 cells (red) on Matrigel embedded with either WT or GαqKO MEFs. F-actin labeling is shown in grey and nuclei in blue. (scale bar 100 µm). Representative images of n=3 independent experiments are shown in all panels. ***, p < 0.001.

To mimic the natural cell environment to a greater extent, we used a three-dimensional (3D)-*in vitro* culture model to evaluate the impact of GFP-expressing WT and GαqKO fibroblasts on red-labeled oral Cal27 tumor cells spheroid formation. Interestingly, we observed that spheroids generated in the presence of GqKO MEFs drastically and significantly increased in number and size compared to their wild-type counterparts (Fig. 2C and Fig. S2B). In addition, the GαqKO-generated spheroids displayed much higher tumor cell spreading and invasion capacity when embedded into Matrigel compared to their WT counterparts (Fig. 2D).

Tumor invasion was further analyzed by confocal microscopy using a 3D organotypic invasion assay. For this purpose, MEFs (either WT or lacking Gαq expression) were embedded into Matrigel, while red fluorescent-labeled tumor cells were seeded on top, mimicking the natural environment of epithelial tissue (scheme in Fig. 2E). Again, the resulting spheroids were significantly larger in the presence of GαqKO compared to WT MEFs. Moreover, only under these conditions did the spheroids gain the ability to enter the Matrigel by following the “railroad” structures generated in the presence of the GαqKO MEFs (Fig. 2E).

Overall, these data strongly suggested a dual role of GαqKO MEFs in promoting tumor growth and invasion, via their higher ECM secretory capacity, the formation of collagen-rich fiber patterns, and additional factors released by these fibroblasts.

### Proteomic analysis reveals an altered endosomal/lysosomal system in GαqKO MEFs

To gain further insight into the molecular mechanisms underlying the tumor-promoting phenotype exerted by GαqKO MEFs, we performed a high-throughput proteomic analysis of these cells compared to the WT control fibroblasts (Supplemental File 1).

Notably, interaction networks and functional annotation enrichment analysis revealed a plethora of biological processes significantly altered in GαqKO MEFs, particularly those related to organelle organization and endocytic trafficking (Fig. 3 A-B) as also evidenced by a completely altered expression profile of most of Rab proteins family members, which are pivotal regulators of both secretory and endocytic pathways as well as of the integrity of intracellular organelles (Fig. 3C). In addition to these changes in intracellular trafficking components, the endocytic machinery itself appeared significantly altered in GαqKO fibroblasts. In the absence of Gαq expression, the clathrin system seems to be downregulated, while a markedly significant upregulation of most of caveolar components (Caveolin1, Caveolin2, Cavin1 and Cavin2) was observed compared to wild-type fibroblasts (Fig. 3C). This proteomic analysis therefore further confirmed our initial observations (Fig. 1A) that Cavin1/PRTF and Cav1 were among the most up-regulated markers in GαqKO fibroblasts.

**Figure 3.**
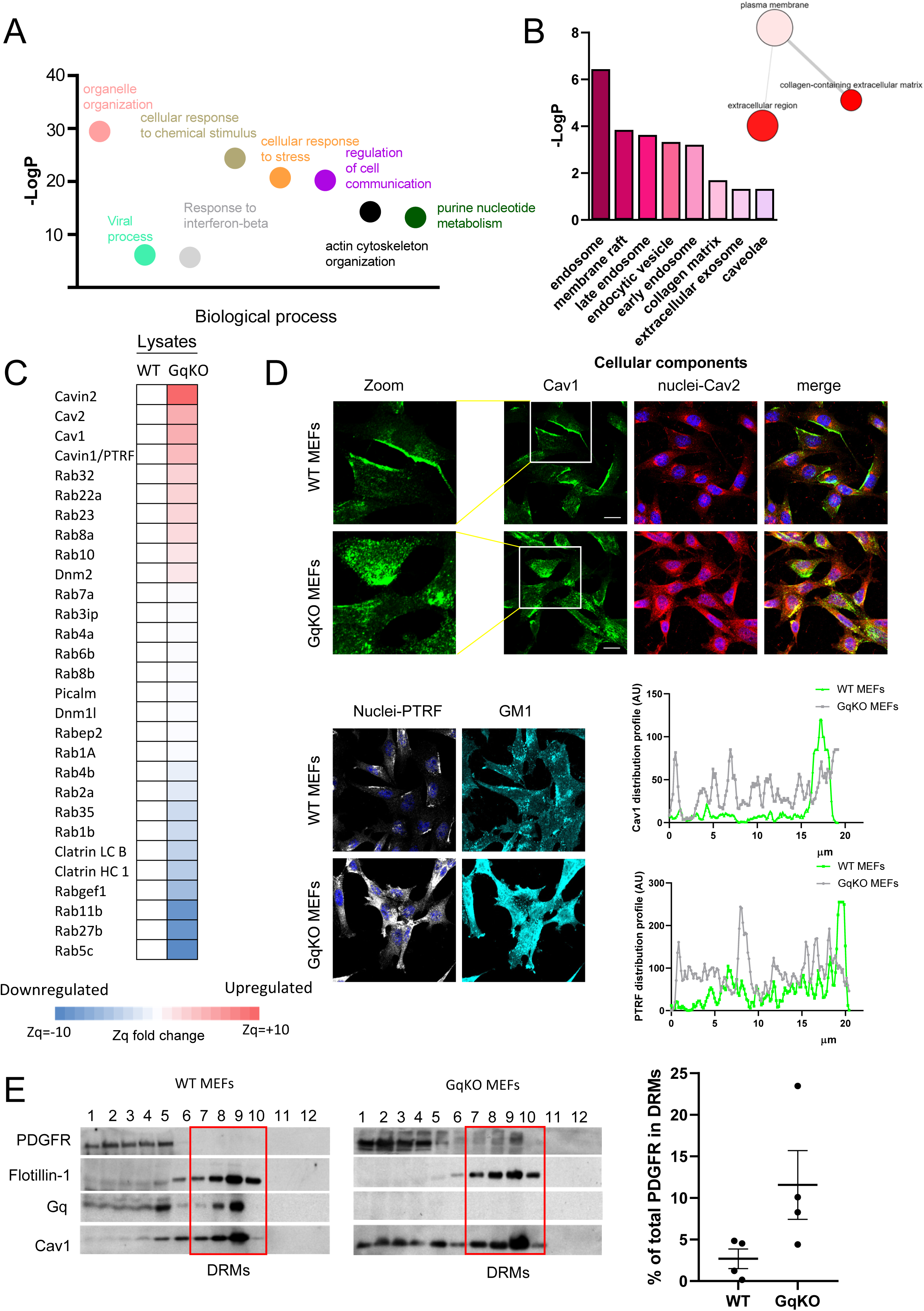
Gαq deficiency in fibroblasts alters the intracellular trafficking networks and the caveolae system. (a) Chart of the most significant GO biological processes that are significantly altered in GαqKO MEFs compared to WTMEFs. Y-axis represents the statistical significance of the enrichment, –log10(p-value). (b) Enrichment diagram of most significant “cellular component” categories upregulated in GαqKO MEFs fibroblasts. Graphic on the right shows a string network corresponding to upregulated extracellular categories. (c) Clustered heatmap of expression of intracellular trafficking proteins altered in GαqKO MEFs compared to WT. The most upregulated family group corresponds to caveolar components. (d) Confocal microscopy analysis of the subcellular distribution of caveolar components caveolin-1 and caveolin-2 (upper panels) and PTRF/cavin and the lipid raft marker GM1 (stained with cholera toxin) (lower panels). Zoom images represent the polarized (WT) versus internalized (GαqKO MEFs) pattern of Caveolin-1. Histograms compares the pixel intensities of caveolin-1 (upper chart) or PTRF (lower chart) in WT and GαqKO MEFs. (e) Western blot analysis of sucrose density gradient fractions from WT and GαqKO MEFs. Fractions were analyzed for the distribution of Cav1, Gαq, Flotillin and PDGFR. Red box denotes detergent-resistant membranes (DRMs) –enriched fractions (7-10). The chart shows the amounts of PDGFR in DRMs relative to the total amount in all fractions; mean±SD (n=4).

To delve into how Gαq deficiency impacts the caveolar system, we examined the subcellular distribution of different caveolar components by confocal microscopy. Our results revealed a clear exclusion of all three analyzed components (Cav1, Cav2, Cavin1/PTRF) from the plasma membrane, along with their accumulation in intracellular membrane structures, thus presenting a phenotype drastically opposite to the preferential membrane localization observed in wild-type fibroblasts (Fig. 3D). In parallel to this altered caveolae distribution, an upregulation of the lipid raft marker GM1 was detected (Fig. 3D), together with a significant enrichment of caveolae as revealed by electron microscopy structures (Fig. S3A). It is noteworthy that subcellular fractionation assays confirmed that caveolar components, regardless of their intracellular compartmentalization, were associated with detergent-resistant membranes (DRMs). Notably, GαqKO fibroblasts showed a clear PDGF receptor (PDGFR) accumulation within these DRM domains that was not detected in WT fibroblasts (Fig. 3E).

Taking together, our data suggested an important involvement of the caveolar system and endocytic trafficking in defining the GqKO phenotype in fibroblasts, with potential implications for key growth factor receptor homeostatic mechanisms.

### Altered PDGFR trafficking and impaired degradation results in its intracellular accumulation in GαqKO MEFs

Given the significantly increased expression levels of PDGFR in GαqKO MEFs (Fig. 1A) and its observed partial enrichment within DRM domains (Fig. 3E), we decided to explore in more detail how the absence of Gαq influences the subcellular localization and trafficking of this receptor. Confocal microscopy studies revealed that, contrary to WT or GαqKI cells, in GαqKO MEFs under basal conditions PDGFR was mainly distributed in intracellular dots (Fig. 4A) that colocalized with Caveolin1 (Fig. S3B) and the lysosomal marker LAMP1 at both low-(Fig. 4B) and high-confluence conditions (Fig. S4A). Such aberrant accumulation and localization of PDGFR in GαqKO MEFs suggested that, although PDGFR was directed to lysosomes for degradation, the final steps of the degradative process failed to function properly. To corroborate this hypothesis, we stimulated WT or GαqKO MEFs with the ligand PDGFβ, a treatment known to induce PDGFR degradation through lysosomal mechanisms ^36^. Upon receptor stimulation, PDGFR was completely degraded in WT MEFs, whereas in GαqKO MEFs, high receptor accumulation was noted, as assessed by both Western blot and confocal microscopy analysis (Fig. 4C), consistent with the notion of impaired PDGFR degradation in such settings.

**Figure 4.**
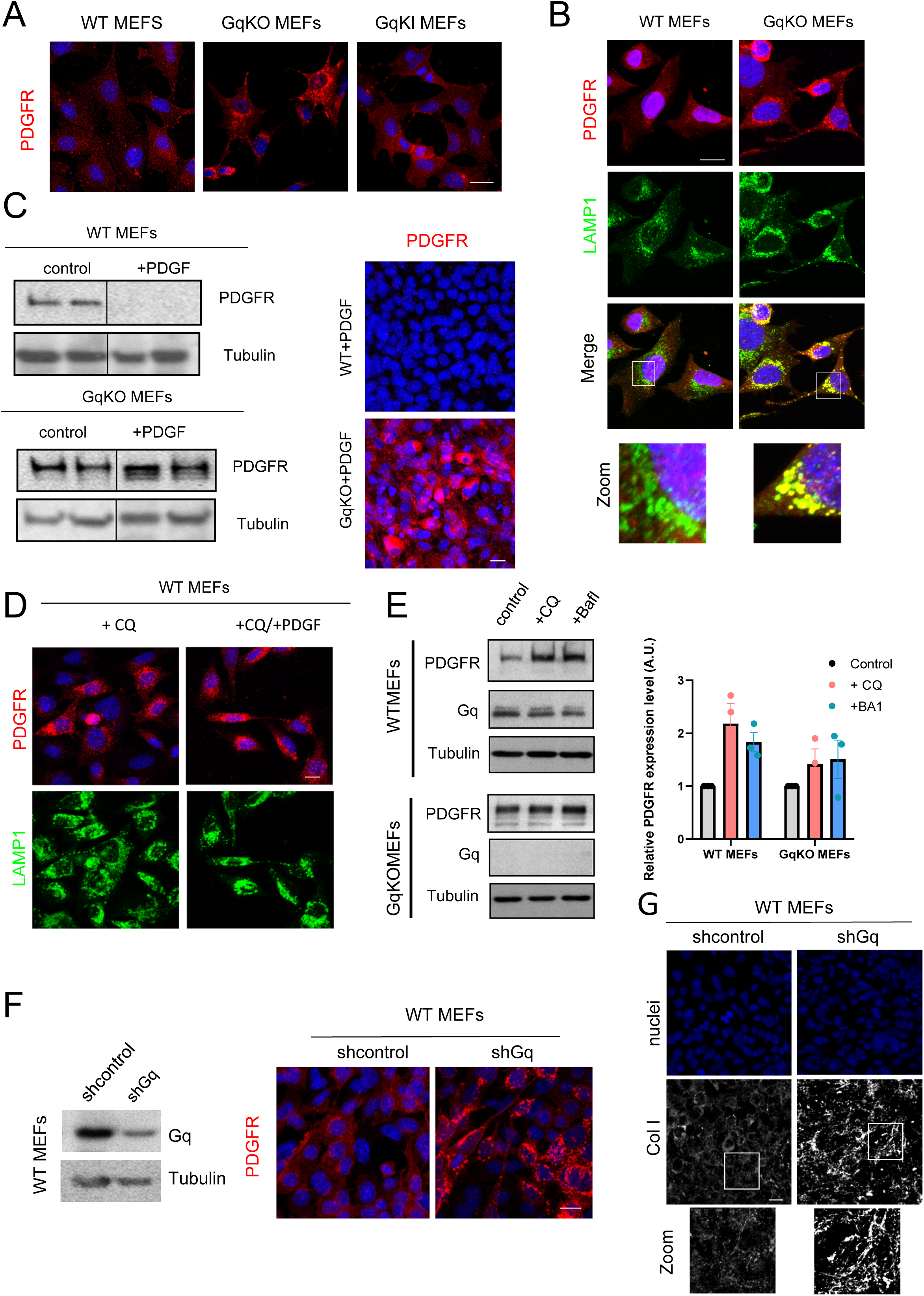
Intracellular accumulation of PDGFR is related to altered trafficking and degradation in GαqKO MEFs. Subcellular localization pattern of PDGF receptors (panels a, b) and of the lysosomal marker LAMP1 (b) in WT, GαqKO and GαqKI MEFs as indicated. Scale bar, 25 µm). (c) Representative western blot (left) and confocal microscopy analysis (right) of PDGFR expression in basal conditions or upon stimulation with PDGF (20 ng/ml) of WT or GαqKO MEFs. Tubulin was used as loading control (scale bar, 25 µm). (d) WT MEFs were treated with Chloroquine (CQ) in the presence or absence of PDGF and analyzed by confocal microscopy using PDGFR (red) and LAMP1 (green) antibodies (scale bar, 10 µm). (e) Western blot analysis of WT and GαqKO MEFs in control conditions or upon treatment with CQ (1µM) or Bafilomycin 1 (BA1 1nM) using PDGFR and Gαq antibodies. Tubulin was used as loading control. Graph represents relative PDGFR levels (mean±SD, n=3). (f) Knock-down efficiency of Gαq in WT MEFs upon lentiviral infection of short-hairpin RNA constructs was assessed by Western blot analysis and PDGFR distribution upon Gαq depletion by confocal microscopy (scale bar, 25 µm). (g) Z-stack projection of confocal microscopy images showing the increased level of collagen I matrix deposition in Gαq-depleted MEFs (scale bar, 25 µm). Zoomed images are also shown.

In this regard, we observed that lysosomal inhibition using either bafilomycin A1 (Baf A1), a drug that impairs lysosomal acidification, or chloroquine (CQ), a classical autophagy inhibitor that prevents autophagosomes from fusing with lysosomes thereby blocking the degradation pathway, led in WT MEFs to increased PDGFR levels, a marked accumulation in intracellular compartments and an enriched distribution in DRMs, even without ligand stimulation (Fig. 4D, 4E and S4B), thus effectively mimicking the phenotype observed in GαqKO MEFs. Notably, GαqKO MEFs PDGFRs remained largely unaffected by these treatments, highlighting their pre-existing defects in lysosomal degradation. In this line, electron microscopy revealed an increased presence of large and fully loaded lysosomal structures in GαqKO MEFs, likely because of altered degradation mechanisms (Fig. S4C).

To confirm the specific dependency of these altered PDGFR modulation patterns on Gαq expression, Gαq knockdown (KD) was performed in WT MEFs through lentiviral infection using a short hairpin RNA targeting this protein. Confocal microscopy analysis revealed that, under these conditions, PDGFR distribution and levels shifted entirely, mimicking the phenotype observed in GαqKO MEFs (Fig. 4F). Interestingly, other hallmark features associated with Gαq deficiency were validated with this silencing approach, including the characteristic increased matrix deposition of collagen I (Fig. 4G), lysosome perinuclear accumulation and the elevated intracellular staining of PTRF/Cavin-1 (Fig. S4D).

Overall, these data strongly indicated that the absence of Gαq expression in fibroblasts alters the degradation efficiency of the lysosomal network, leading to enhanced PDGFR, levels and marked intracellular accumulation.

### Exosomes represent the preferred route of PDGFR release in the absence of Gαq expression

To explore how GαqKO MEFs can maintain homeostasis despite their lysosomal dysfunction, we evaluated the contribution of exosome release as a potential adaptive strategy to reduce cellular stress and to partially maintain the levels of proteins that cannot be efficiently degraded at the lysosomal level. Increasing evidence demonstrates that autophagic and exosomal pathways are interconnected ^23,37^, and our previous studies have shown that Gαq plays a crucial role in controlling autophagy ^31^ which may influence exosome release through autophagy-dependent secretion mechanisms. Thus, we chose to investigate whether PDGFR could be sorted and released through the exosomal pathway in GαqKO MEFs as a compensatory mechanism.

In general, the process of protein release through exosome generation requires its prior sorting into multivesicular bodies (MVB). We confirmed that in GαqKO MEFs the internal pool of PDGFR strongly colocalized with the MVB marker LBPA (Fig. 5A). Notably, strong clusters of PDGFR localized to intraluminal vesicles (ILVs) within MVBs were also clearly observed when endosomes were artificially enlarged by expressing the constitutively active form of Rab5 (Rab5Q67L) (Fig. S5A). These observations suggested that MVBs are one of the destinations of endocytosed PDGFR in GαqKO MEFs.

**Figure 5.**
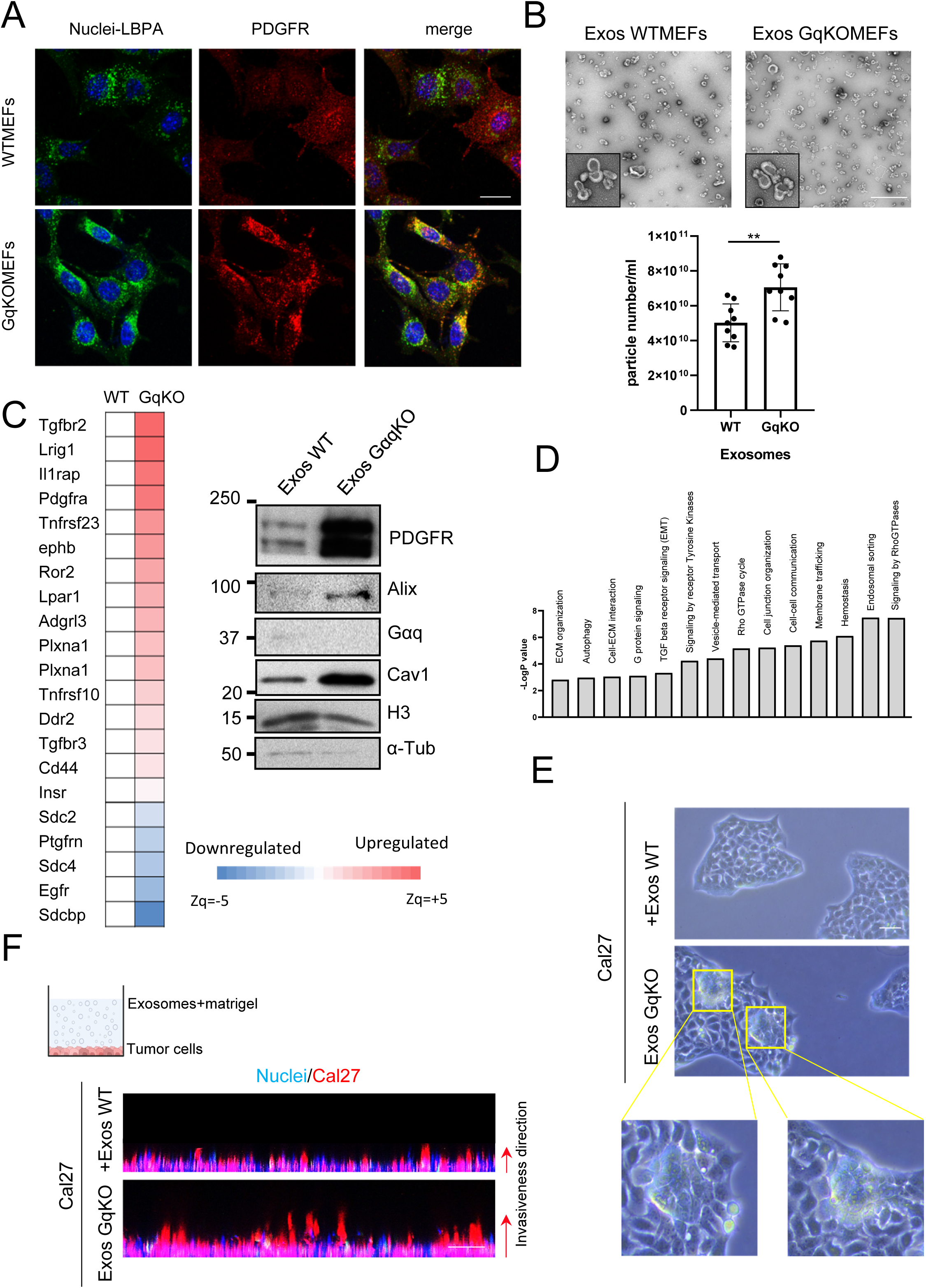
GαqKO MEFs secrete exosomes aberrantly loaded with PDGFR and other factors that promote oral cancer cells tumoral features. (a) Confocal microscopy analysis of the subcellular distribution of the MBV marker LBPA (green) and PDGFR (red) in WT and GαqKO MEFs. Colocalization is shown in yellow (scale bar, 25 µm). (b) Exosomes were isolated from culture supernatants of WT and GαqKO fibroblasts. Representative electron microscopy images (scale bar, 200 nm) and quantification of exosome particles relative to cell number are shown (mean±SD; n= 8). (c) List of the most significantly up-or down-regulated growth factor receptors present in GαqKO fibroblasts-derived exosomes, and Western blots analysis of the indicated proteins in exosomes derived from equal cell numbers of WT and GαqKO MEFs. (d) Enrichment diagram of “cellular component” category for proteins upregulated in GαqKO MEF– derived exosomes compared to WT. (e) Phase-contrast microscopy showing morphological changes in human oral cancer Cal27 cells 48 h after exposure to exosomes derived from either WT or GαqKO MEFs (scale bar, 50 µm). Zoomed images point to aberrant tumor growth features. (f) Representative Matrigel invasion assay of Cal27 tumor cells through Matrigel containing WT or GαqKO MEFs-derived exosomes. A scheme of the protocol is shown in the upper., **, p < 0.01.

To evaluate whether this accumulation of PDGFR may led to its specific release via exosomes, we purified and analyzed exosomes produced by either WT or GαqKO MEFs. Interestingly, in the absence of Gαq fibroblasts exhibited a slight but consistent increase in the number of exosomes released (Fig. 5B). More strikingly, exosomes derived from GαqKO fibroblasts exhibited a drastic and substantial enrichment of PDGFR compared to those released by their wild-type counterparts (Fig. 5C) along with other markers such as Alix or Cav1. This phenotype was mimicked when WT MEFs were treated with the lysosomal inhibitor CQ (Fig. S5B), consistent with the notion that a disrupted degradative mechanism in GαqKO MEFs redirects PDGFR towards exosome-mediated secretion.

To further explore the differential exosome sorting modulated by Gαq expression, we performed a comparative proteomic analysis of the exosomes released by WT and GαqKO MEFs (Supplemental file 2). This analysis not only confirmed the strong and significant upregulation of PDGFR within the exosomes produced by GαqKO MEFs, as previously observed by Western blot analysis, but also intriguingly uncovered that a large number of other specific tumor growth factor receptors were markedly present in the exosomes derived from GαqKO (Fig. 5C and Supplemental file 2), along with other changes in components of secretory, trafficking and signaling networks (Fig. 5D) Therefore, in GαqKO MEFs, exosome release appears to represent an alternative pathway to facilitate the secretion of many different tumor growth factors otherwise unable to undergo proper lysosomal degradation.

In this regard, we compared the influence of exosomes purified from either WT or GαqKO MEFs on 2D cultures of oral cancer Cal27 cells. After 48 hours, Cal27 cells exposed to GαqKO-derived exosomes showed and enhanced pattern of tumor growth compared to their controls (Fig. 5E). Moreover, these cells also displayed an enhanced invasive capacity, as assessed by a 3D-invasion assay (Fig. 5F). These data indicated that exosomes derived from GαqKO MEFs display a pronounced tumor-promoting effect, consistent with the substantial presence of growth factor receptors within these vesicles.

### Aberrant desmoplastic stroma-bearing tumors are induced by GαqKO MEFs in vivo

To validate the role of GαqKO MEFs in orchestrating a more favorable tumor-promoting environment compared to their wild-type counterparts *in vivo*, orthotopic tongue allografts were generated by injecting luminescent and GFP-expressing Cal27 cells, either alone or in combination with WT or GαqKO MEFs, in immunodeficient NSG mice, followed by tumor tracking by bioluminescence-based imaging (Fig. 6). Strikingly, seven days after injection, animals co-injected with GαqKO MEFs exhibited rapid and pronounced tumor growth at the tongue, as detected by bioluminescence imaging (Fig. 6A), macroscopically (Fig. S6A) and by confocal microscopy analysis of histological tumor tissues (Fig. 6C) leading to a severe mice condition, which forced us to euthanize the animals approximately 10-12 days after cell injection following the established humane endpoint protocol (Fig. 6B). Interestingly, when we used Gαq-reconstituted GαqKO fibroblasts (GαqKI), survival and tumor progression data were like those observed in mice co-injected with WT MEFs (Fig S6B), underscoring the critical role of stromal Gαq in mitigating tumor-promoting effects.

**Figure 6.**
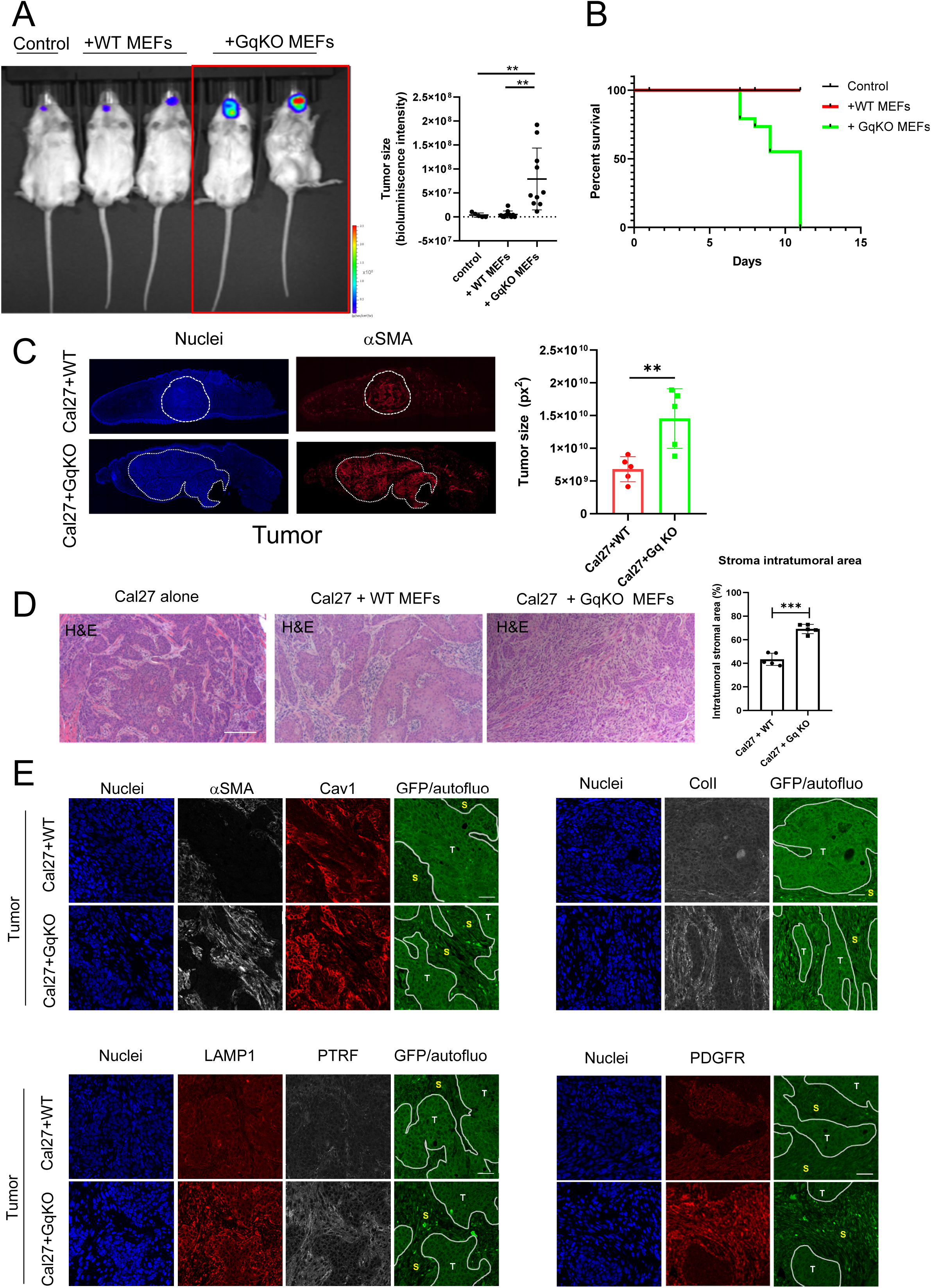
GαqKO MEFs trigger desmoplastic and aggressive tumors in vivo. (a) Bioluminiscent imaging of orthotopic tongue allografts. in mice transplanted with a luciferase-expressing tumor tongue cancer cell line (Cal27-luc) either alone (control) or in combination with WT or GqKO MEFs. Color scale correlates with luminescence count production. The graph displays the quantification of tumor cell-derived bioluminescence (n=10 mice per condition, One-way ANOVA). (b) Survival curve for control animals (tumor cells alone) and those co-injected with WT or GαqKO MEFs. (c) Representative tissue sections of complete tongue tumors generated as in panel a, showing nuclei (blue) and αSMA (red) staining. The chart represents tumor size (mean±SD, n=5 mice/group) in the indicated conditions. (d) Hematoxylin-Eosin staining at 10X magnification of tumor samples corresponding to the different conditions, obtained after eleven days of cancer cells injection into the mice. The graph illustrates the amount of stroma in tumors generated under each condition (mean±SD, n=5 mice/group). (e) Immunohistochemical staining patterns in the tumor samples of the different proteins markers indicated. S, stroma; T, tumor areas, delineated by dotted lines. **, p < 0.01; ***, p < 0.001.

Remarkably, histological analysis of the generated tumors by hematoxylin/eosin staining revealed that tumors formed in the presence of GαqKO fibroblasts exhibited a pronounced stromal desmoplastic reaction, a phenotype that was markedly different from that observed in tumors generated by Cal27 cells alone or co-injected with WT MEFs (Fig. 6D), stressing the relevance of GαqKO fibroblasts in determining tumor microenvironment features.

A detailed immunofluorescence analysis of the tumors generated *in vivo* confirmed a robust stromal upregulation of key markers also detected in *in vitro* settings, such as αSMA, Cav1, PDGFR, LAMP1 or PTRF, specifically in the tumors generated by co-injection with GαqKO MEFs (Fig. 6E). Regarding matrix organization, collagen I analysis revealed a stronger and much stiffer collagen matrix surrounding tumors in conditions involving GαqKO MEFs, as determined through immunostaining (Fig. 6E) and multiphoton excitation microscopy coupled with second harmonic generation (MPE-SHG) imaging (Fig S6C). Taken together, our results strongly indicated a critical role of GαqKO MEFs in driving aberrant tumor growth, stromal remodeling, and increased tumor aggressiveness *in vivo*.

### Human-derived HNSCC CAFs exhibit features similar to those observed in GαqKO fibroblasts

In HNSCCs, a heterogeneous CAFs population plays a pivotal role in accelerating tumor progression and invasion by secreting growth factors, remodeling the ECM, altering vasculature, and increasing therapy resistance ^38,39^.

We thus explored whether Gαq expression is altered in human HNSCC-derived CAFs, by Western blot analysis of CAF subpopulations obtained from minced human tumor tissue of surgically resected HNSCC and of Normal fibroblasts (NFs) obtained simultaneously from healthy tissue of the same patients (n of 6, Fig. 7A). Although variability was observed between sample pairs, we found a moderate downregulation of Gαq expression in the CAFs compared to NFs, which correlated with an upregulated PDGFR expression in such tumor fibroblasts. In addition, CAFs showed a more spindle-like morphology compared to NFs (Fig. 7B), which was associated with a high intracellular accumulation of PDGFR (Fig. 7C), enhanced F-actin and αSMA labeling (Fig. 7D), and a marked deposition of ECM components such as FN and Col I (Fig. 7E). All these features were like those displayed by GαqKO MEFs and were further recapitulated upon specific Gαq knockdown (KD) performed on human normal fibroblasts (NFs) through lentiviral infection using the previously described short hairpin RNA targeting this protein (Fig. S7A and S7B).

**Figure 7.**
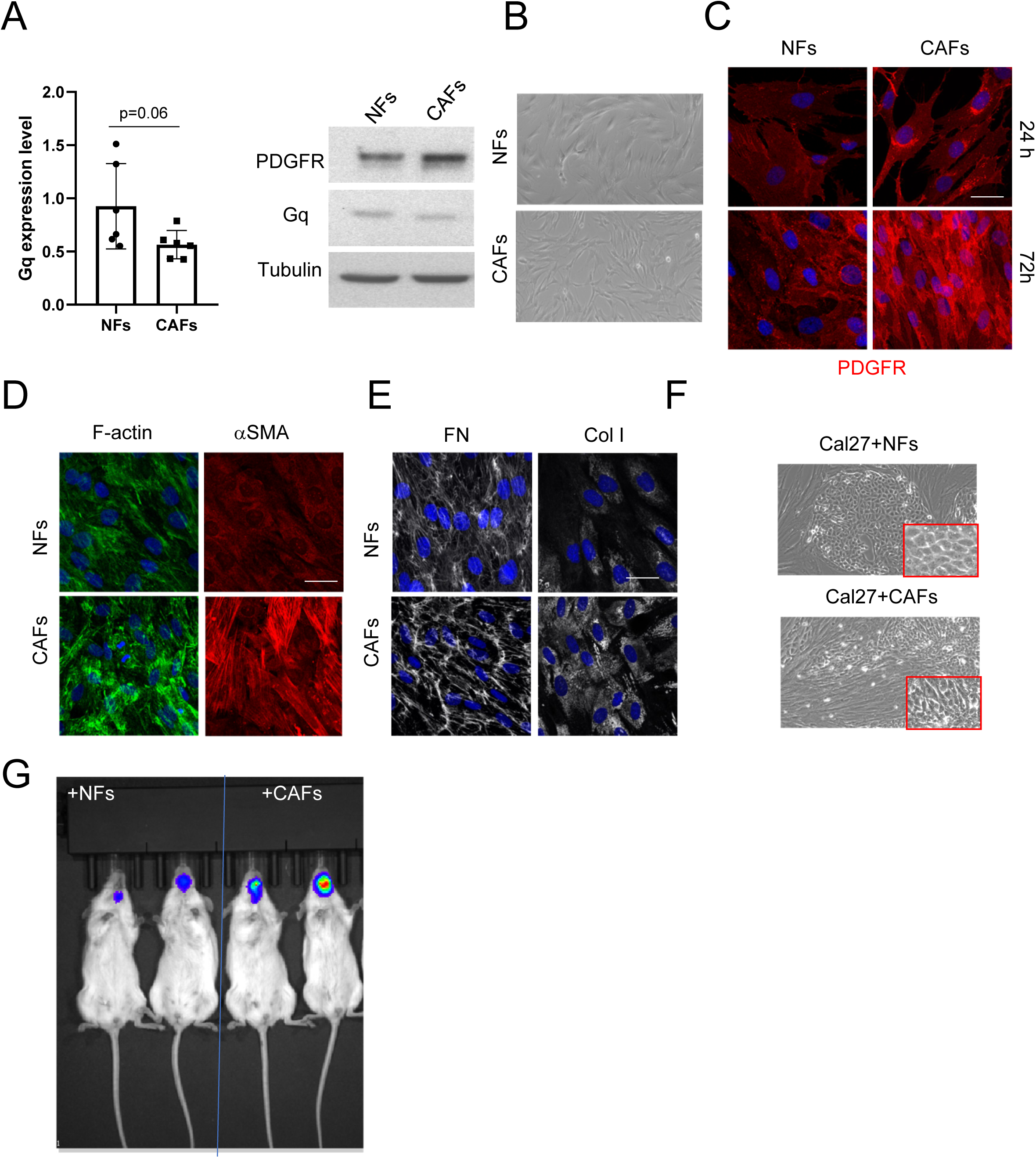
Patient-derived HNSCC CAFs display reduced Gαq expression and foster enhanced tumor progression of oral cancer cells. (a) Western blot analysis and quantification of Gαq expression in Normal Fibroblasts (NFs) and Cancer-Associated-Fibroblasts (CAFs) derived from HNSCC tumor patients (mean±SD, n=6 patients). PDGFR expression levels were also measured, using Tubulin as a loading control. (b) Phase contrast microscope images of one selected pair of NF-CAFs. (c) PDGFR subcellular distribution in the selected NF-CAF sample at either low or high confluency (scale bar 25 µm). (d) Confocal microscopy images displaying F-actin (green) and α-SMA (red) staining of NFs and CAFs. (e) Z stack projection of confocal microscopy images showing Fibronectin (left panels) and Collagen I (right panels) in matrix deposited by NFs or CAFs (scale bar, 25 µm). (f) Phase contrast images of a 2D coculture of the Cal27 human oral cancer cells and either NFs or CAFs. Zoomed images illustrate differences in tumor cell distribution and cell-cell contact patterns. (g) Bioluminescence imaging of orthotopic tongue allografts in mice transplanted with a luciferase-expressing tumor tongue cancer cell line (Cal27-luc) in combination with NFs or CAFs. Color scale correlates with luminescence count production.

Interestingly, the 2D coculture of Cal27 oral cancer cells with patient-derived CAFs resulted in a phenotype characterized by a disorganized and more chaotic organization and reduced tumor cell-cell contact, similar to features observed upon coculture with GαqKO MEFs. In contrast, the presence of NFs induced the formation of a highly confined and encapsulated tumor cell distribution (Fig. 7F). *In vivo*, all these traits manifested as a notably aggressive and invasive tumor phenotype upon orthotopic tongue co-injection of bioluminescent Cal27 cells with CAFs, again resembling features of GαqKO MEFs, whereas tumors formed with NFs exhibited a more contained and less invasive profile (Fig. 7G). Overall, our data provide compelling evidence that stromal Gαq expression plays a pivotal role in shaping the tumor microenvironment and in controlling HNSCC tumor progression.

## Discussion

HNSCC is one of the most severe and challenging types of cancer due to its high level of heterogeneity and the limited availability of efficient therapeutic strategies ^10^. Continuous co-evolution and communication of tumor cells with the TME seems to be determinant in HNSCC progression ^40^. Cancer-Associated Fibroblasts (CAFs) are key components of the TME, and essential in shaping tumor dynamics and driving HNSCC progression. Therefore, a better comprehension of the mechanisms that govern CAFs features and their interactions with other TME components is crucial for developing targeted therapies that could improve treatment outcomes for HNSCC patients^10,41,42^.

We unveil here an unforeseen pivotal role of fibroblast Gαq in fostering HNSCC cancer progression by remodeling the tumor microenvironment (TME). MEFs lacking Gαq exhibit elevated expression of specific fibroblast activation markers and enhanced matrix remodeling and contractile features, resembling those of tumor-promoting CAFs. Additionally, Gαq-deficient fibroblasts exhibit altered endocytic and degradative machinery, resulting in aberrant trafficking and impaired degradation of PDGFR, a critical driver of HNSCC progression. This dysfunction enhances the selective cargo sorting of various growth factor receptors, including PDGFR, into released exosomes. Functionally, GαqKO MEFs sustain oral HNSCC tumor cell growth and invasiveness in *in vitro* assays and markedly promote the formation of desmoplastic stroma-bearing tumors *in vivo*. Furthermore, most of these phenotypic traits are also observed in patient-derived HNSCC CAFs, which display a reduction in Gαq expression levels (Figure 8).

**Figure 8.**
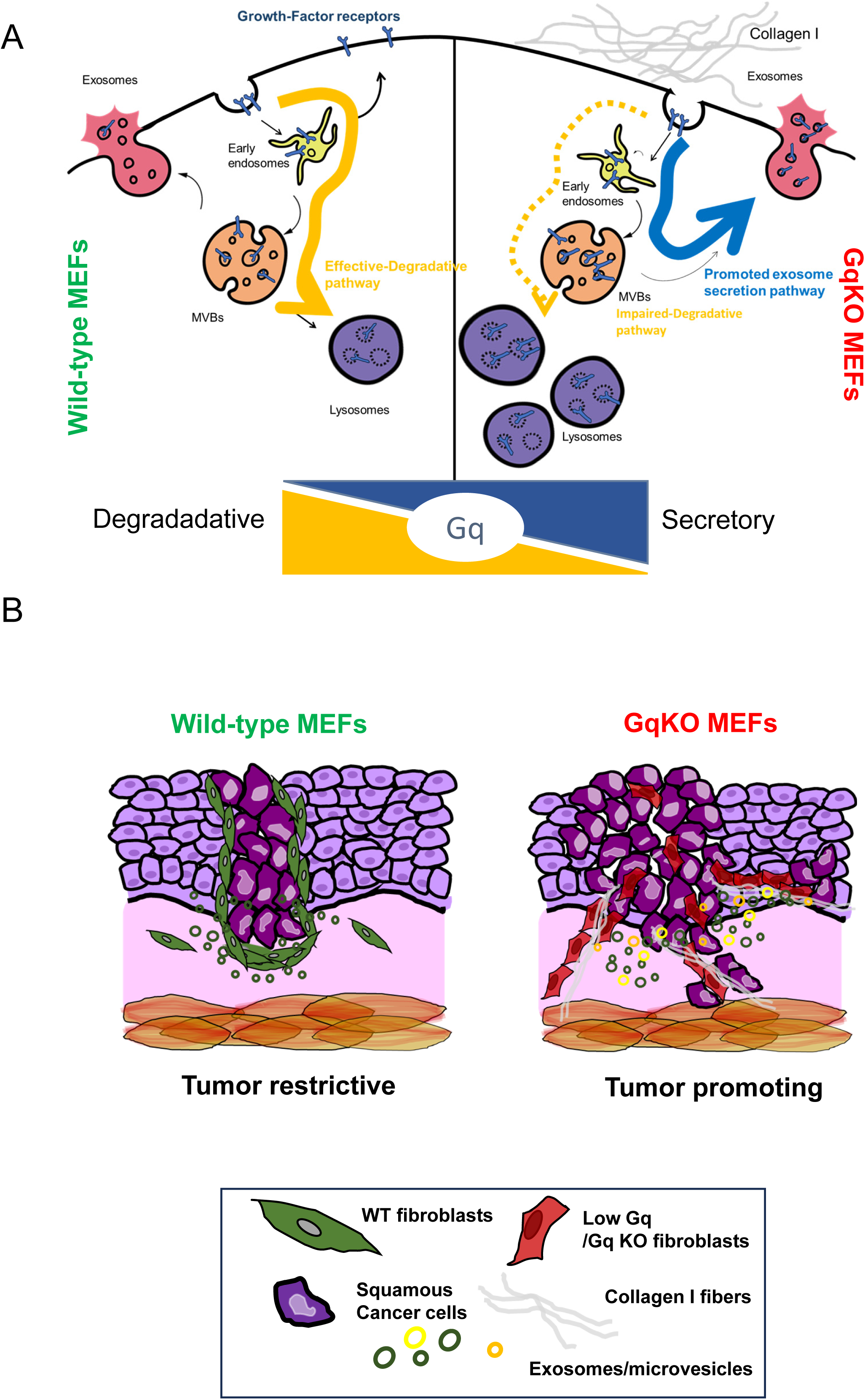
Proposed model for the role of stromal Gαq in HNSCC tumor progression. Gαq expression in the tumor stroma emerges as a key regulator of fibroblast traits and ofin HNSCC progression. (a) The absence of Gαq expression in fibroblasts switches trafficking and degradative pathways towards a preferentially secretory phenotype. This context favors the specific sorting of tumor growth factor receptors and their release via the exosomal pathway, rather than their degradation taking place in wild-type mouse embryonic fibroblasts (WT MEFs). Such altered secretory capacity is also reflected by significant collagen I matrix deposition in GqKO MEFs. (b) At the functional level, a collagen I-rich matrix and the secretion of exosomes highly enriched in tumor growth factor receptors driven by GqKO downregulation fosters aberrant growth and dissemination of oral cancer cells.

In general, ECM organization is a key determinant in tumor aggressiveness ^43,44^. In HNSCC, a strong desmoplastic reaction driven by CAFs enhances invasiveness ^5^. These cells exhibit diverse subpopulations with both tumor-promoting and tumor-suppressing roles, depending on their interaction with the ECM ^45–47^. Notably, CAFs that disrupt the ECM-epithelial boundary promote invasion, while those forming defined barriers hinder it, underscoring the complexity of CAF behavior in cancer. In this context, Gαq-deficient fibroblasts display a pro-tumorigenic phenotype, characterized by a collagen-rich, linearized matrix with a chaotic distribution that significantly promotes oral HNSCC cell invasion.

Regarding CAFs markers, α-SMA is particularly important in HNSCC, where its expression levels have been shown to correlate with patient outcomes ^10,48^. Prolonged culture and confluence drive GαqKO MEFs toward a myofibroblast-like phenotype, characterized by a marked increase in α-SMA expression, enhanced contractility, and augmented ECM remodeling, including organized collagen I fibers. Interestingly, our proteomic analysis also reveals that GαqKO MEFs can exhibit traits of inflammatory fibroblasts, such as high expression of PLA2, CD34, and JAK/STAT pathways. Thus, our results suggest the possibility that GqKO MEFs may transition from pro-inflammatory to a myofibroblastic status overtime. This aligns with the intrinsic plasticity of CAFs, which can dynamically shift between myofibroblastic, inflammatory, and antigen-presenting phenotypes in response to cues from tumor milieu, as evidenced by recent single-cell RNA sequencing studies ^49–51^. Further investigation will be required to confirm this switch.

PDGFR also serves as a key biomarker for the identification of CAFs in oral squamous cell carcinoma ^52^. A striking feature linked to Gαq deficiency in fibroblast is the marked alteration of PDGFR trafficking and degradation. This results in its intracellular accumulation and enhanced release within exosomes, along with a variety of growth factor receptors. Recent research has emphasized the potential of CAF-secreted exosomes as important prognostic makers^53,54^. However, despite the promising role of CAF-secreted exosomes as potential targets for cancer, the mechanisms governing their biogenesis and secretion remain poorly understood. One possibility relies on the crosstalk of exosomes biogenesis with the autophagy process^23^. Our group has previously described that Gαq-deficient MEFs exhibit increased autophagy both in basal and under nutrient stress conditions^31,55^, which may progressively evolve from a primarily degradative process to a secretory phenotype over time. This shift is evidenced by the marked accumulation of lysosomes in the perinuclear region in GαqKO-MEFs^31^, thus influencing their activity^56^ and favoring the redirection of proteins targeted for degradation towards exosomal secretion, as we describe herein for the PDGFR and other receptors.

It is thus tempting to suggest that an altered balance of the endolysosomal trafficking and degradative networks takes place upon Gαq downregulation, promoting pro-tumoral changes in exosome cargo sorting and secretion. Consistent with this notion, we detect marked changes in the dosage of key components of the trafficking (Rab proteins networks) and endocytic (Caveolin-1 and related proteins) machinery in GαqKO MEFs compared to their WT counterparts.

Rab GTPases are key determinants of organelle identity within the endomembrane system, orchestrating vesicle tethering and membrane fusion to ensure proper intracellular trafficking, also playing a vital role in exosome biogenesis ^57–59^. Our proteomic analysis reveals significant alterations in a variety of components of the Rab machinery in Gαq-deficient MEFs, which would likely contribute to the rerouting of vesicular cargo and to the observed unbalance between autophagic degradation and exosomal release.

Interestingly, caveolae internalization and trafficking is also directly dependent on Rab proteins modulation^60^. The caveolae system is crucial for the interaction between membrane organization and the regulation of GPCRs and tumor growth factor receptors, influencing key cellular processes such as signaling and receptor turnover ^61–64^. Strikingly, our proteomic study reveals that in GαqKO MEFs the clathrin-mediated internalization pathway is downregulated, while we detect a strong upregulation and altered distribution of caveolar components such as the key core molecule caveolin-1 (Cav-1), accompanied by increased intracellular accumulation and a loss of polarized plasma membrane distribution. Such “internalization” of Cav-1 from its usual plasma membrane localization seems to correlate with an enhanced exosome release in GαqKO MEFs. Interestingly, Cav-1 has recently emerged as a new regulator of exosome biogenesis and protein cargo sorting via both conventional exosome pathways and autophagy-dependent mechanisms ^65,66^. On the note, Cav-1 modulates PDGFR signaling by directly binding to PDGF receptors and inhibiting their function. When PDGFR dissociates from Cav-1, it undergoes clathrin-dependent internalization and is targeted to lysosomal degradation ^63,67–69^. We observe in GαqKO fibroblasts a redirection of PDGFR from lysosomal degradation to exosomal sorting, which correlates with increased PDGFR internalization into Cav-1 positive compartments. Overall, our data support the notion that the absence of Gαq shifts intracellular trafficking of PDGFR and other growth factor receptors from a clathrin-dependent to a caveolae-mediated internalization process, thereby favoring secretion over degradation (Scheme in Fig.8).

Changes in lipid composition may also play a critical role in intracellular organelle trafficking and may help to explain some of the phenotypes observed in GαqKO MEFs. Effective lipid segregation is essential for determining the composition and fate of intracellular vesicles^70,71^. Interestingly, Gαq is a key regulator of cellular lipids, as its activation initiates the phospholipase Cβ (PLCβ) signaling cascade, leading to the cleavage of phosphatidylinositol 4,5-bisphosphate (PIP2) to diacylglycerol (DAG) and inositol 1,4,5-trisphosphate (IP3)^72^. Indeed, Gαq activation is also required to activate PLCβ isoforms depending on βγ released by Gi/o and/or Gs heterotrimeric complexes^73^. It is tempting to suggest that the absence of Gαq may disrupt this lipid balance, resulting in an altered lipid profile in fibroblasts that may contribute to the observed increase in caveolae formation and changes in intracellular vesicles dynamics. Thus, further research into the functional links between Gαq, Caveolin-1, Rab GTPases and lipid modulation could shed light on the CAF-like features triggered upon Gαq downregulation.

Importantly, we find that cancer-associated fibroblasts from HNSCC patients show a slight reduction in Gαq expression and exhibit similar features to those observed in GαqKO MEFs, including enhanced PDGFR expression and intracellular accumulation, and increased collagen I deposition. Moreover, silencing Gαq in normal human fibroblasts triggers the CAF-like features of GαqKO MEFs. Functionally, our data show that the altered remodeling of the extracellular matrix (ECM) and the secretory features associated with reduced Gαq expression in both human HNSCC CAFs and GαqKO MEFs can promote the growth and invasiveness of oral HNSCC tumor cells in 2D and 3D co-culture assays *in vitro* and strongly promote the formation of desmoplastic stroma-bearing tumors *in vivo*, suggesting a central role for Gαq function in the tumor stroma of HNSCC patients (Fig. 8).

In sum, our data strongly suggest that altered Gαq expression in the tumor stroma acts as a novel and critical regulator of HNSCC progression through a dual mechanism: (1) promotion of a collagen I-rich matrix that facilitates tumor dissemination and (2) secretion of exosomes highly enriched in tumor growth factor receptors because of an imbalanced degradative capacity that drives aberrant tumor growth. A better understanding of how cancer-associated fibroblasts integrate the modulation of the ECM architecture and metabolic properties (via autophagy/degradation and exosome secretion pathways) in a Gαq-dependent manner may help to design novel therapeutic strategies targeting the TME to halt HNSCC progression.

## Materials and methods

### Reagents and plasmids

The following primary antibodies were used: anti-PTRF (ab48824), TSG101 (AB83), PDGFR (ab32570) and GNAQ (ab75825) were from Abcam. Anti-tubulin (T-6074), anti-Fibronectin (F3648), anti-Smooth muscle actin (A5228) and FSP1 (ABF32) were from Sigma. Rabbit monoclonal caveolin1 (D46G3), Histone 3 (9717S), Alix (3A9) and E-Cadherin (4065) were from Cell Signaling; Rat LAMP1 (1D4B) LAMP-1) was from Hybridoma Bank Alexa fluor 488, 546 and 647-conjugated phalloidin and fluorescent secondary antibodies were from Life Technologies. Matrigel (354230) and Collagen I from rat tail (354249) were from Corning. Bafilomycin (1nM, Sigma-Aldrich), Chloroquine (1 µM, Sigma-Aldrich), or a specific Gαq/11/14 inhibitor (YM-254890, 5 µM, Wako) was used as indicated in the figure legends. For generation of enlarged endosomes/MVBs, transfection of cells with the plasmid expressing the constitutive active form of Rab5 (Rab5(Q79L)) was performed.

### Animal model

NOD.Cg-Prkdcscid Il2rgtm1Wjl/SzJ (NOD-SCID IL2rγnull 4; NSG) immunodeficient mice were kindly provided by Dr. Miguel R. Campanero García from Centro de Biología Molecular Severo Ochoa (CBM). All mice were bred under specific pathogen-free conditions in accordance with CBM institutional guidelines. Experiments were performed in accordance with Spanish legislation on animal protection and were approved by the local governmental animal care committee (Proex 118.8-21). HNSCC cancer cell injections in the tongue were conducted on young mice ranging from 9-14 weeks old. Implications of gender was also considered including both males and females as part of our studies.

### Cell Culture

Wild-type and Gαq/11-Knock-Out (referred to in the manuscript as GαqKO) Mouse embryonic fibroblasts (MEFs) were kindly provided by Dr. S. Offermanns (Max-Planck-Institute for Heart and Lung Research, Germany). These MEFs were generated using the Cre/loxP recombination system in floxed alleles of the Gq and G11 genes. MEFs were cultured in Dulbeccós modified Eagle medium (DMEM) supplemented with 10% fetal bovine serum (FBS), 100 U/ml penicillin and 100 µg/ml streptomycin at 37°C in a humidified 5% CO2-95% air incubator.

UMSCC-47 and HN13 and Cal27 (RRID:CVCL_1107), (kindly provided by Dr J.S Gutkind UCSD, USA), oral cancer cells were cultured in DMEM/F-12 supplemented with 10% FBS, 100 U/ml penicillin and 100 µg/ml streptomycin.

Normal fibroblasts (NFs) and cancer-associated fibroblasts (CAFs) were kindly provided by Dr. Juana García Pedrero and Dr. Saúl Alvarez Teijeiro (Department of Otolaryngology and Instituto Universitario de Oncología del Principado de Asturias, University of Oviedo, Spain). Primary cancer-associated fibroblasts (CAFs) were obtained from minced tumor tissue of surgically resected HNSCC patients at the Hospital Universitario Central de Asturias. Normal dermal fibroblasts (NF) were obtained from the dermis of human foreskin. They were cultured in DMEM supplemented with 10% fetal bovine serum (FBS) (Gibco, Waltham, MA, USA), 100 U/mL penicillin 200 mg/mL streptomycin, 2 mM L-glutamine, 20 mM HEPES (pH 7.3) and 100 mM MEM non-essential amino acids. All cell lines were tested periodically for mycoplasma contamination by PCR.

### Generation of Lentiviral vectors and reconstituted GαqKO MEFs cell lines

A pSD44-Gαq wild-type vector (Addgene) was used for lentiviral reconstitution of GαqKO fibroblasts and generation of the GαqKI MEFs cell line.

For knocking down Gαq experiments, Gαq was silenced by infection of cells with shRNA vectors; used sequence targeting human/mouse Gαq (sequence CACCAAGCTGGTGTATCAGAA) was cloned into the short hairpin lentiviral vector pLVX-shRNA2, which contains a ZsGreen1 reporter (Clontech), and the vector was used to transfect HEK293T cells. Cells for experiments were exposed to HEK293T cell supernatants and infection efficiency was monitored by Zsgreen 1 expression.

### Generation of Luciferase-expressing oral cancer cell lines

Cal27 cells were transduced with lentiviral particles generated with the pIRSIN-Luc-IRES-GFPfp vector, a generous gift from Patricia Fuentes (Centro de Biología Molecular, Spain) by incubation with 8 μg/ml Polybrene. GFP (and Luciferase)-positive cells were isolated by cell sorting using a FACSAria Fusion system in the Flow cytometry service of the Centro de Biología Molecular Severo Ochoa.

### Orthotopic injection of cancer cells and fibroblasts

A total volume of 50 μl was injected into the tongue of 9-14-week-old NSG mice containing either 1×10^6^ Cal27 cancer cells or 1×10^6^ Cal27 cancer cells + 1×10^6^ of the indicated MEFs variants to induce orthotopic tumors. The Cal27 cell line stably expresses GFP and luciferase. In the experiments performed using NFs and CAFs, the injected volumes contained either 750.000 Cal27 cancer cells or 750.000 Cal27 cancer cells+750.000 NFs or CAFs. Orthotopic tumors were analyzed by immunohistochemistry (see below) and by bioluminescence acquisition from anesthetized mice (isofluorane gas 1.5%, Abbott, Madrid, Spain) inoculated in each condition using an IVIS Spectrum after intraperitoneal injection of 150 mg/kg of body weight of D-Luciferin (Promega, Madison, WI), according to the manufacturer’s instructions.

### Histological analysis and immunohistochemistry

Tissue samples were fixed in 10% neutral buffered formalin (NBF) overnight and embedded in paraffin wax. Sections (5 μm) were stained with H&E or processed for immunohistochemistry. Briefly, first step was deparaffinization of tissue sections by 15-min incubation at 55°C and later submersion, 3 times in Histoclear and 3 times in 100% ethanol, followed by 95, 70 and 50% ethanol and distilled water washes. Antigen retrieval was carried out in citrate buffer pH 6 for 5 min in the microwave and cooling down 15 min at room temperature. After washing with PBS twice (5 min each wash), sections were then incubated with blocking solution (3% Horse-Serum-PBS) for 1 h. Primary antibody dilution was made in blocking solution, and the incubation was carried out in the humidity chamber overnight at 4°C. Sections were stained with antibodies against FN, Collagen I, α-SMA, Ki67, Caveolin-1, PDGFR, PTRF and E-Cadherin. The following day, sections were washed extensively twice with PBS and once with 0.1% Tween-PBS. Secondary antibody conjugated with specific fluorophores were diluted in blocking solution and incubated for 2 h (1:200). Sections were washed twice with PBS and once with 0.1% Tween-PBS. Finally, the tissue sections were mounted on cover slides and analyzed by confocal microscopy.

### Immunofluorescence microscopy of cultured cells

Cells grown on glass coverslips were washed twice with PBS and fixed for 15 min at room temperature in 4% paraformaldehyde in PBS. Fixed cells were washed extensively with PBS and permeabilized for 5 min in PBS containing 0.1% Triton X100 and 2% BSA to reduce non-specific binding. Cells were incubated for 1 h at room temperature in PBS containing 0.2% BSA with primary antibodies (typically diluted 1:200). Subsequent washes and incubations with Alexa fluor-647 or 546 phalloidin or fluorescent secondary antibodies were carried out in PBS for 1 h at room temperature. Coverslips were mounted in Permafluor aqueous mounting medium and examined with a Laser Scanning Confocal Microscope LSM800 coupled to an inverted Axio Observer Microscope (Zeiss) typically through 40X (ACS APO 40.0x 1.15 OIL) or 63X (ACS APO 63.0x 1.30 OIL) objectives.

### FN fiber quantification

The procedure for image processing and quantification of FN length fiber size and thickness extracellular was done using a macro in Fiji (ImageJ 1.50e x64) as described in Albacete-Albacete et al., 2020^65^

### Western blot and cell fractionation

Cell proteins were resolved by SDS-PAGE and transferred to nitrocellulose membranes. Blots were incubated for 2 h in TBS containing 5% bovine serum albumin and overnight with primary antibodies (typically diluted 1:1000). After incubation (2 h) with horseradish peroxidase conjugated goat anti-rabbit or goat anti-mouse antibody, signal on washed blots was detected by enhanced chemiluminescence (GE Healthcare). Band intensities were quantified with Image Lab (Bio-Rad).

### Preparation of detergent-resistant membrane fractions

Detergent-Resistant-Membranes (DRMs) derived from WT or GqKO MEFs lysates were purified as described in Navarro-Lerida et al. JCS (2002)^74^. Briefly, two p150 plates of each fibroblast cell line were scraped and resuspended in 2 ml (final volume) of MES buffered saline (MBS: 25 mM MES, pH 6.5, 0.15 M NaCl, 1 mM PMSF plus 1% Triton X-100) at 4°C. Cells were homogenized by a minimum of 10 strokes through a syringe (0.5×16 mm) on ice. The homogenate was brought up to 4 ml by adding 2 ml 80% sucrose in MBS and placed at the bottom of a Beckman SW40 13 ml Ultraclear tube. The discontinuous sucrose gradient (40-30-5%) was formed by sequentially loading 4 ml 30% sucrose and 4 ml 5% sucrose in MBS. Cells fractions were separated by centrifugation at 200,000 g for 18 h in a SW40 rotor (Beckman) at 4°C. A light, scattered band confined to the 5-30% sucrose interface was observed that contained most caveolin-1 protein and excluded most other proteins. Twelve (1 ml) fractions were collected, starting from the bottom of the tube. An equal volume of cold acetone was added to each fraction and the proteins precipitated overnight at 4°C. Protein pellets were collected by centrifugation at 16,000 g and air dried for 2 h to eliminate acetone traces. The protein precipitates were then analyzed by SDS-PAGE and western blot.

### Spheroid formation assay

After trypsinization, Cal27 cells were resuspended in FBS-free DMEMF12, counted, and adjusted to the desired concentration (usually 1X10^4^ cells/ml). A 20 µL final volume was prepared with 5X methyl cellulose in FBS-free medium and the cancer cell suspension. The mixture was carefully deposited in a drop on the inner surface of a 150 mm dish lid. The lid was placed on a plate containing 10 ml PBS to humidify the culture chamber. Under gravity, cells aggregated at the bottom of the hanging drop. After 24 hours, the resulting cell aggregates were lifted with a pipette. For 3D spheroid invasion assays, the aggregates were embedded into 25-μl 1:1 mix of Matrigel and medium or, alternatively, spheroids were deposited onto a monolayer generated by WT or GαqKO fibroblasts. After 4-6 days, fixation with PFA 4% in PBS was performed and immunofluorescence and confocal microscopy was used to monitor cell migration/invasion in each condition.

### Collagen Gel contraction assay

The assay was performed as previously described^75^. Briefly, 2X10^5^ MEF cells of the different genotypes were mixed with NaOH-titrated collagen I (Corning) to a final collagen I concentration of 1 mg/ml. The mixture was immediately transferred to a 24-well plate and lattices were allowed to solidify. Serum-containing medium was added to each well and gels were manually detached by circular movements using a sterile pipette tip. Gels were placed at 37°C, and contraction was documented.

### Isolation and characterization of exosomes released by fibroblasts

Exosomes were isolated from cultured fibroblasts grown in exosome-free culture medium. To remove detached cells, conditioned medium was collected and centrifuged at 300 g for 10 min at 4°C. The supernatant was collected and centrifuged at 2,000 g for 20 min at 4°C. The supernatant was then centrifuged at 10,000 g at 4°C for 30 min to completely remove contaminating apoptotic bodies, microvesicles, and cell debris. The clarified medium was then ultracentrifuged at 110,000 g at 4°C for 70 min to pellet the exosomes. The supernatant was carefully removed, and crude exosome-containing pellets were washed in ice-cold PBS. After a second round of ultracentrifugation, the resulting exosome pellets were resuspended in the desired volume of PBS. The protein precipitates were monitored by Western blot for the expression of exosomal markers Alix or Tsg101. In all experiments, the exosomes used corresponded to the total exosome pellets resulting from the serial centrifugation steps; usually, 10^6^ exosome particles were added per cell. Exosome uptake was performed in exosome-free medium. Exosome concentrations and size distributions were determined by Nanoparticle Tracking Analysis (Nanosight).

### Electron Microscopy

Exosomes were visualized by TEM according to the method of Thery et al. (2006)^76^. The exosome suspension was fixed in 2% PFA and transferred to formvar/carbon–coated EM grids. After 20 min, grids were placed sample-face down for 2 min in a 100-μl drop of PBS on a sheet of parafilm. Grids were then transferred to 1% glutaraldehyde for 5 min and then washed with distilled water. Samples were contrasted in 2% uranyl acetate and examined by TEM.

In addition, WT and GqKO MEFs were processed for EM following standard procedures. Briefly, cells were fixed in 0.1 M cacodylate buffer, pH 7.4 containing 2.5% glutaraldehyde (+1 mg/ml ruthenium red), and then post-fixed in 1% osmium tetroxide (+1 mg/ml ruthenium red), followed by treatment with 2% uranyl acetate. The samples were dehydrated, embedded in LX112 Epon resin, sectioned and stained.

### Second Harmonic generation microscopy

SHG imaging on in vivo tumor generated by orthotopic injection of cancer cells and fibroblasts in the tongue was done as described^77^

### Proteomic analysis

Lysates derived from WT or GqαKO MEFs, or exosomes derived from these cell lines were digested using the filter aided sample preparation (FASP) protocol^78^. Briefly, samples were dissolved in 50 mM Tris-HCl pH 8.5, 4% SDS and 5 mM Tris(2-carboxyethyl) phosphine (TCEP), boiled for 10 min, and centrifuged. Protein concentration in the supernatant was measured with the Direct Detect® Spectrometer (Millipore). About 100 μg of protein were diluted in 8 M urea in 0.1 M Tris-HCl pH 8.5 (UA buffer) containing 5mM TCEP and loaded onto 30 kDa centrifugal filter devices (Millipore). The reduction buffer was removed by washing with UA, and proteins were then alkylated by incubation in 20 mM iodoacetamide in UA for 20 min in the dark. Alkylation reagents were eliminated by washing three times with UA and three additional times with 50 mM ammonium bicarbonate. Proteins were digested overnight at 37°C with modified trypsin (Promega) in 50 mM ammonium bicarbonate at a 40:1 protein:trypsin (w/w) ratio. The resulting peptides were eluted by centrifugation with 50 mM ammonium bicarbonate (twice) and 0.5M sodium chloride. Trifluoroacetic acid (TFA) was added to a final concentration of 1% and the peptides were finally desalted onto Oasis-HLB cartridges and dried-down for further analysis.

For the quantitative analysis, tryptic peptides were dissolved in 150 mM triethylammonium bicarbonate (TEAB) buffer, and the peptide concentration was determined by measuring amide bonds with the Direct Detect system. Equal amounts of each peptide sample were labelled using the 10plex TMT Multiplex reagents (Thermo Fisher). Briefly, each peptide solution was independently labelled at room temperature for 1 h with one TMT reagent vial previously reconstituted with acetonitrile (ACN). After incubation at room temperature for 1 h, the reaction was stopped with diluted TFA and peptides were combined. Samples were concentrated in a Speed Vac, desalted onto Oasis-HLB cartridges, and dried-down for mass spectrometry analysis.

Digested peptides were loaded into the LC-MS/MS system for on-line desalting onto C18 cartridges and analyzed by LC-MS/MS using a C-18 reversed phase nano-column (75 µm I.D. x 50 cm, 2 µm particle size, Acclaim PepMap RSLC, 100 C18; Thermo Fisher Scientific) in a continuous acetonitrile gradient consisting of 0-30% B for 360 min and 50-90% B for 3 min (A= 0.1% formic acid; B=100% acetonitrile, 0.1% formic acid). A flow rate of 200 nl/min was used to elute peptides from the RP nano-column to an emitter nanospray needle for real time ionization and peptide fragmentation in an Orbitrap Fusion mass spectrometer (Thermo Fisher). During the chromatography run, we examined an enhanced FT-resolution spectrum (resolution=70,000) followed by the HCD MS/MS spectra from the nth-most intense parent ions. Dynamic exclusion was set at 40 s. For increased proteome coverage, labelled samples were also fractioned by high-pH reverse phase chromatography using C18 microcartridges (Thermo Fisher). Fractions were analyzed using the same system and conditions described before.

Protein identification and quantification. All spectra were analysed with Proteome Discoverer (version 2.1.0.81, Thermo Fisher Scientific) using SEQUEST-HT (Thermo Fisher Scientific). The Uniprot database, containing all mouse sequences (March 03, 2013) concatenated with decoy sequences generated using DecoyPyrat^79^, was searched with the following parameters: trypsin digestion with 2 maximum missed cleavage sites; precursor and fragment mass tolerances of 2 Da and 0.03 Da, respectively; methionine oxidation as a dynamic modification; and carbamidomethyl cysteine and N-terminal and Lys TMT6plex modifications as fixed modifications.

For quantitative analysis, the iSanXoT program ^80^ quantifies the intensity of TMT reporter ions derived from the isobaric labeling of fragmentation spectra. First, the false discovery rate (FDR) was calculated using the corrected Xcorr score (cXcorr)^81,82^, with an additional filter for precursor mass tolerance of 12 ppm^83^. Identified peptides had an FDR of 1% or lower.

Next, peptide-to-protein assignment was performed using an in-house developed method, where each peptide is assigned to the most likely protein in the UniProtKB/Swiss-Prot database. Proteins are ranked according to the number of peptides with which they are identified. The algorithm assigns each peptide to the protein with the highest number of peptides. In cases of tied proteins, it prioritizes assignments based on the number of peptide-spectrum matches.

Subsequently, quantitative information from TMT reporter intensities was integrated from the spectrum level to the peptide level, and then to the protein level, to quantify the relative abundance of each protein. This was done according to the WSPP (weighted spectrum, peptide, and protein) statistical model ^84^ and the Generic Integration Algorithm (GIA) ^85^. This model standardizes peptide and protein abundance quantification as log2-ratios, with values expressed in units of standard deviation for peptides (Zp) and proteins (Zq) according to their estimated variances.

Differences in protein abundance or functional behavior were estimated by comparing the groups’ Zq or Zc medians, respectively, as determined by the WSPP statistical model. Proteins or functional changes were considered statistically significant with a two-sided t-test comparison p-value < 0.05 of the Z values.

Functional class enrichment was assessed by GSEA using the IPA™ (Qiagen) and DAVID resources; gene subsets related to ECM remodeling were further manually curated from the MSigDB hallmark gene set collection 75. Visualization of enriched functional annotation terms was performed using the open source REVIGO suite and or GOrilla software. The mass spectrometry proteomics data have been deposited to the ProteomeXchange Consortium via the PRIDE ^86^ partner repository with the dataset identifier **PXD061641**.

### Statistical analysis

Error bars depict SEM or SD as indicated in figure legends. Statistical significance was determined with GraphPad Prism by unpaired Student’s t-test for one-to-one comparisons or by 2-way-ANOVA in the case of multiple comparisons.; * p<0.05, ** p<0.01, *** p<0.001.

## Supporting information

Supplementary infomation Navarro-Lerida et al.,

## ACKNOWLEGMENTS

We thank Dr. J.S. Gutkind (UCSD, USA), S. Offermanns (Max Planck, Germany) and Patricia Fuente (CBM, Spain) for experimental tools, and Pilar Hernández and Angustias Page (CIEMAT, Spain) for histologic processing of the samples. We thank CNIC Proteomics Unit and CBMSO Animal Care, Advanced Light microscopy, Electron Microscopy, Flow cytometry and Preclinical Biomedicine facilites for their technical assistance and support. We would also like to thank Dr. Miguel R. Campanero and Dr. Alberto Hernández Alcántara (CBMSO) for their generous provision of NSG mice.

This work was supported by Instituto de Salud Carlos III (grants PI22_00966 to C-R, PI22/00167 to JMG-P and PI24/00398 to SA-T, co-funded with European FEDER contribution), Agencia Estatal de Investigación of Spain (grant PID2023-146735OB-I00 to F-M), H2020-MSCA Program, Grant agreement 860229-ONCORNET 2.0 to F-M), CIBERCV-Instituto de Salud Carlos III, Spain (grant CB16/11/00278 to FM, co-funded with European FEDER contribution), Fundación Ramón Areces grants (to F-M and C-R), Programa de Actividades en Biomedicina de la Comunidad de Madrid-S2022/BMD-7209 –INTEGRAMUNE (to F-M.). CIBERONC--Instituto de Salud Carlos III (CB16/12/00390 to JMG-P, co-funded with European FEDER contribution). LP-F and RH-L are recipients of a FPU predoctoral fellowship from the Spanish Ministry of Education (FPU20/01588 and FPU22/00979 respectively) and SA-T is a recipient of a Miguel Servet research contract from ISCIII (CP23/00101) and co-funded with European FEDER contribution. We also acknowledge institutional support to the CBM from Fundación Ramón Areces.

The CNIC is supported by the Instituto de Salud Carlos III (ISCIII), the Ministerio de Ciencia, Innovación Universidades (MICIU) and the Pro CNIC Foundation), and is a Severo Ochoa Center of Excellence (grant CEX2020-001041-S funded by MICIU/AEI/10.13039/501100011033).

## Author contributions

IN-L provided the study of concepts, designed and performed experiments, prepared figures, analyzed data and wrote and revised manuscript; RH-L, GP-G, MIJ-L performed some experiments, analyzed data and prepared figures; DG-M assisted with animal experiments. JA-L performed proteomic analysis. MAdP contributed to the study concepts, LP-F, SA-T and JMG-P provided human NFs and CAFs, and contributed to the study concepts; FM Jr, and CR contributed the study of concepts, wrote and revised manuscript and obtained funding.

