## Supplementary infomation Navarro-Lerida et al., for "Loss of Gαq reshapes key fibroblast traits and drives matrix remodeling and aggressive progression of oral cancer tumors"

#### Supplementary Figure Legends

**Figure S1. Lack and inhibition of Gαq activity modify fibroblast-specific features.** (a) Phase contrast microscope images of WT and GαqKO MEFs after 24 h of seeding (20x magnification). (b) Confocal analysis of F-actin and Gαq subcellular distribution in WT and GαqKO MEFs (scale bar 10 µm). (c) Western blot expression of Cav1 and PDGFR in fibroblasts after 96 h of culture. (d) Fibronectin matrix deposition (in grey) and lysosomal distribution (in red) in WT MEFs treated with the Gαq inhibitor YM254890 (5µM). A LAMP1 antibody was used as lysosome marker (scale bar 10 µm).

**Figure S2. Effect of co-cultured fibroblasts on Cal27 oral cancer cells.** (a) Microscopy images of co-cultures of Cal27 cells with WT or GαqKO fibroblasts. Cells were stained for the epithelial-mesenchymal transition marker E-cadherin (in grey), αSMA (in green) and F-actin (in green); nuclei were stained with Hoechst. Zoomed images show E-cadherin redistribution from cell-cell contact sites to intracellular compartments in the presence of KO fibroblasts (arrows) (scale bar, 10 µm). (b) Phase contrast analysis of spheroids formed in Matrigel co-culture of Cal27 tumor cells and either WT or GαqKO MEFs for 72 h.

**Figure S3. Altered trafficking and degradation of PDGFR in GαqKO MEFs**. (a) WT and GαqKO MEFs were analyzed by electron microscopy. A higher number of caveolae (indicated by arrows) are present in GαqKO MEFs compared to WT (scale bar 100 nm). (b) Colocalization of PDGFR (green) and Caveolin-1 (red) in GαqKO MEFs as determined by confocal microscopy. Merged yellow signals are better observed in the zoomed area.

**Figure S4. Altered PDGFR/lysosomal pattern associated with Gq deficiency in fibroblasts**

(a) Confocal microscopy analysis of LAMP1 (green) and PDGFR (red) subcellular distribution in WT and GαqKO MEFs under confluent conditions (scale bar, 25 µm). (b) PDGFR expression distribution by Western blot analysis of sucrose density gradient fractions of WT and GαqKO MEFs and WT MEFs treated with Bafilomycin1 (1nM). Red box denotes DRMs-enriched fractions. (c) Electron microscopy image of an aberrantly loaded lysosome in GαqKO MEFs. Scale bar 200 nm. (d) Distribution pattern of F-actin, LAMP1 and PTRF upon Gαq silencing in WT MEFs by lentiviral infection with a short hairpin RNA targeting Gαq.

**Figure S5. Enhanced PDGFR trafficking into multivesicular bodies (MVBs) leading to exosome secretion in GαqKO MEFs**. (a) Distribution of PDGFR (red) in Rab5(Q79L)-expressing MEFs (grey). Aberrant accumulation of PDFGR in MVBs is observed preferentially in GαqKO MEFs. (b) Western blots analysis of the indicated proteins in exosomes derived from PDGF-treated WT cells in the absence or presence of chloroquine (CQ, 1 μM). TSG101 is used as an exosomal marker.

**Figure S6. Large desmoplastic stroma-bearing tumors are induced by GαqKO MEFs in vivo.** (a) Representative picture of macroscopic tongue tumor growth upon orthotopic allografts generated by injection of Cal27 oral cancer cells alone or in combination with WT or GαqKO MEFs. (b) Bioluminescence detection of tumor cells generated by orthotopic injection of Cal27 cells alone or in combination with either GαqKO MEFs or a Gαq-reconstituted version (GαqKI MEFs) (n=4 animals per condition). (c) Representative images of self-assembled collagen matrix in the indicated tumors, as measured by second harmonic generation microscopy.

**Figure S7. Effects of Gαq knockdown on normal human fibroblasts.** (a) Knock-down efficiency of Gαq upon lentiviral infection of short-hairpin RNA constructs in human normal fibroblasts obtained from tongue healthy tissue was assessed by Western blot analysis, along with levels of PDGFR, Cav1 and GAPDH as loading control. Quantification of Gαq levels is shown in the graph (mean± SD, n=3). (b) Confocal images showing the effect of Gαq silencing on LAMP distribution (red, zoomed area is included) and on FN matrix deposition (grey). Scale bar, 50 μm). **, $p < 0.01$.

### Figure S1

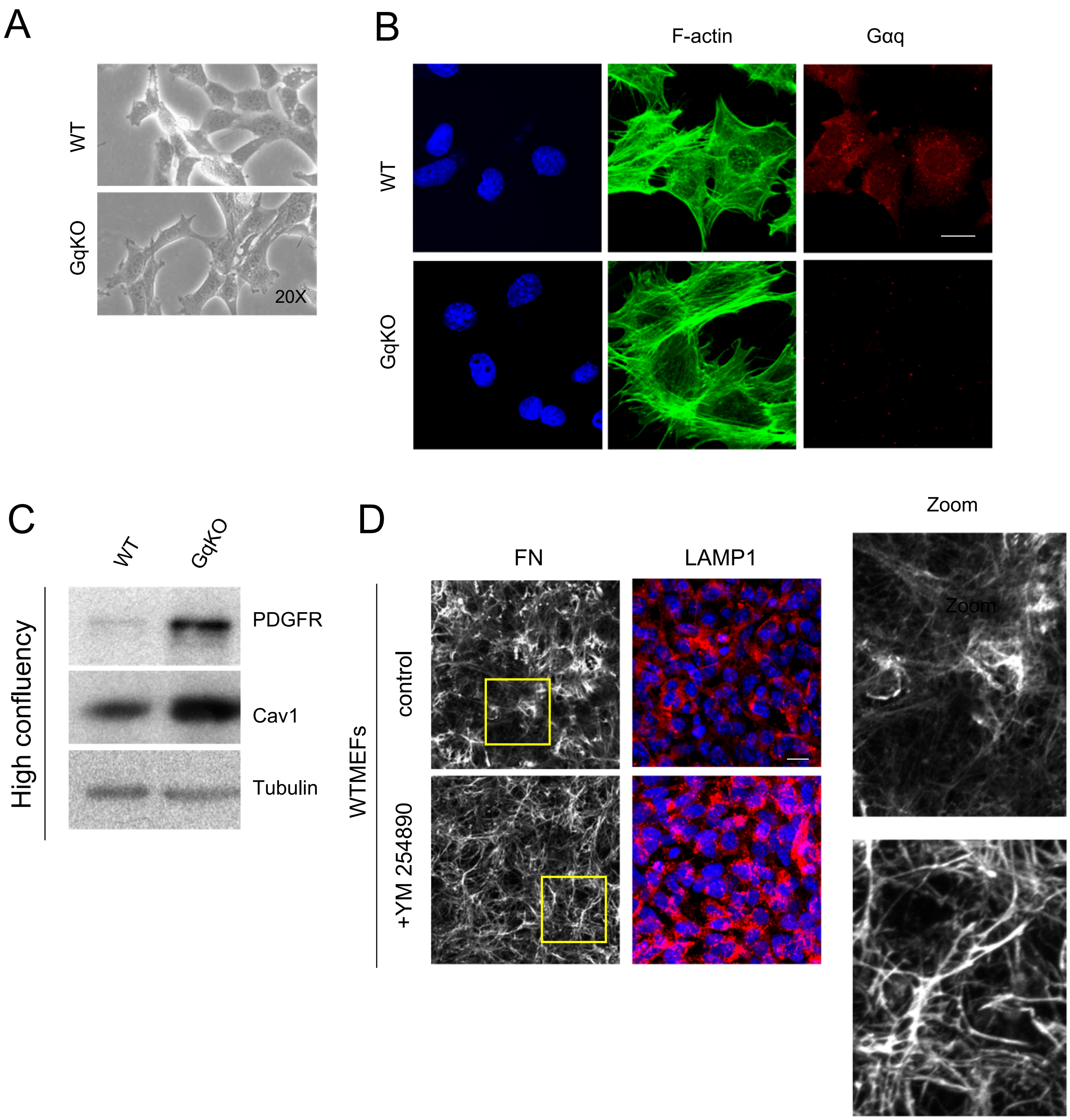

A
WT
GqKO
20X
B
F-actin
Gαq
WT
GqKO
C
WT
GqKO
High confluency
PDGFR
Cav1
Tubulin
D
FN
LAMP1
Zoom
WTMEFs
control
+YM 254890
Zoom

### Figure S2

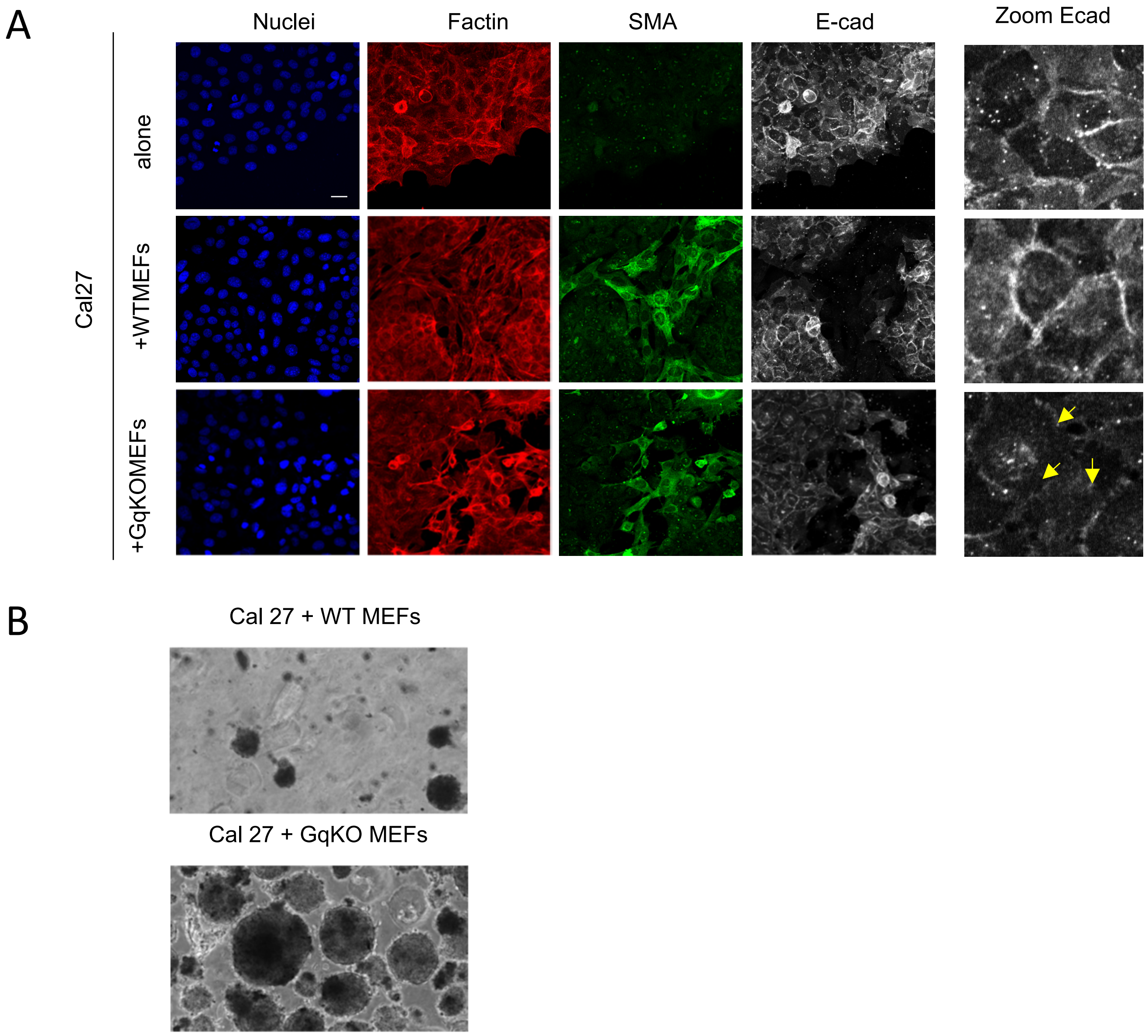
A
Nuclei
Factin
SMA
E-cad
Zoom Ecad
Cal27
alone
+WTMEFs
+GqKOMEFs
B
Cal 27 + WT MEFs
Cal 27 + GqKO MEFs

### Figure S3

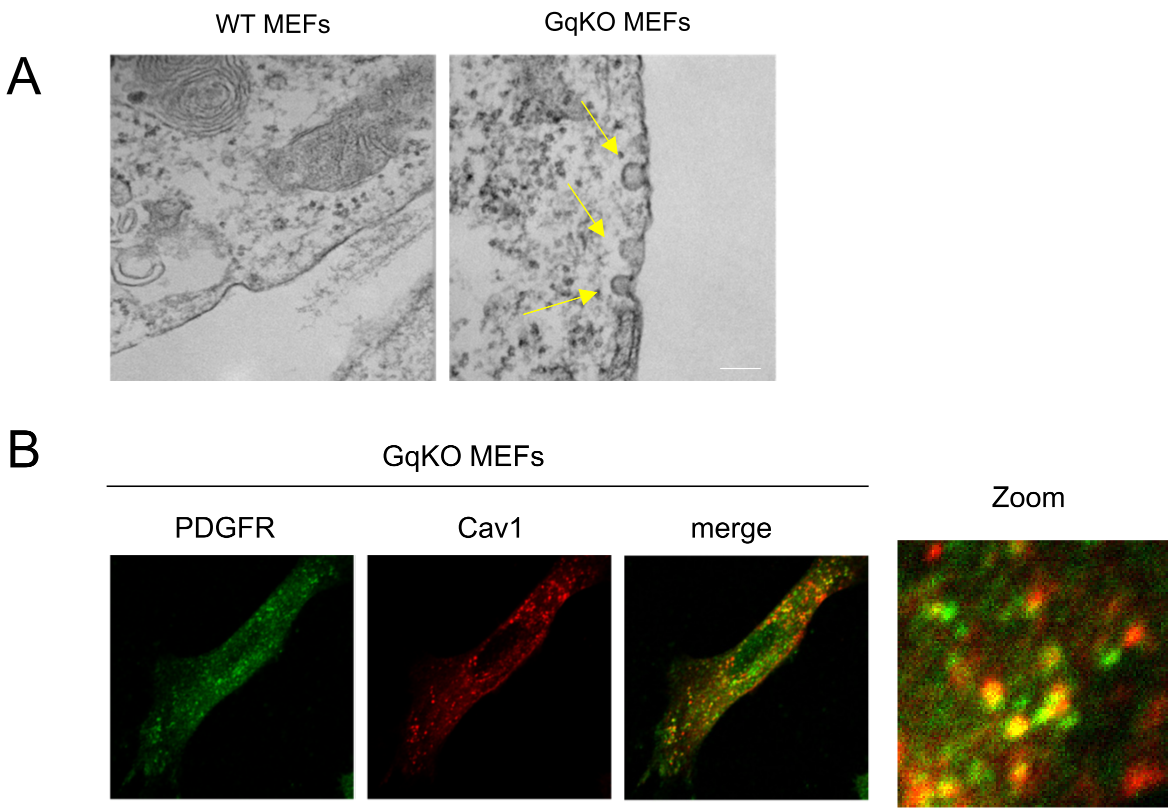

### Figure S4

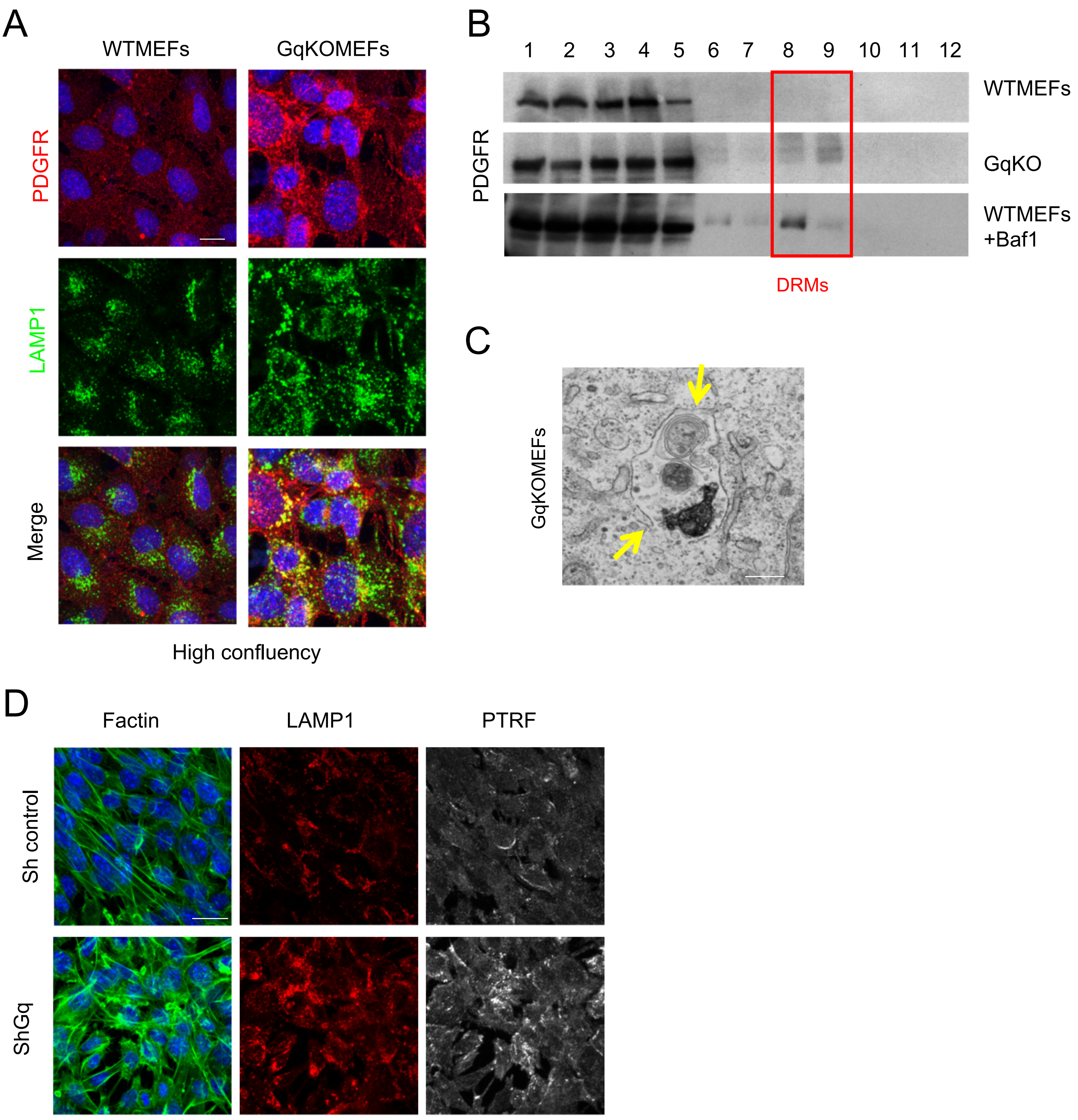
A
WTMEFs
GqKOMEFs
PDGFR
LAMP1
Merge
High confluency
B
1
2
3
4
5
6
7
8
9
10
11
12
PDGFR
WTMEFs
GqKO
WTMEFs
+Baf1
DRMs
C
GqKOMEFs
D
Factin
LAMP1
PTRF
Sh control
ShGq

### Figure S5

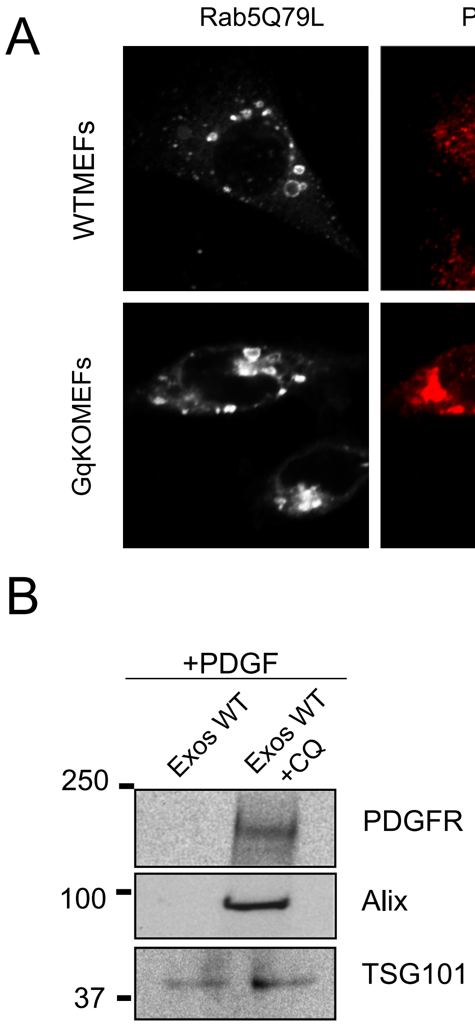
A
Rab5Q79L
PDGFR
merge
WTMEFs
GqKOMEFs
B
+PDGF
Exos WT
Exos WT +CQ
250
100
37
PDGFR
Alix
TSG101

### Figure S6

A

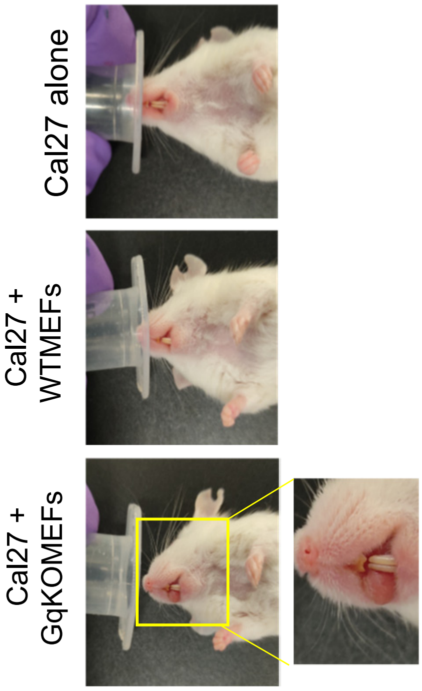


B

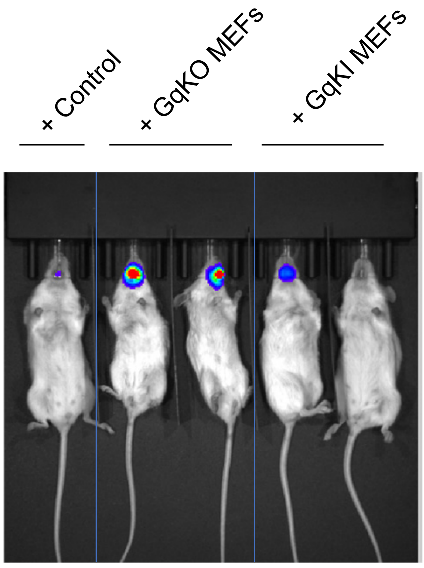


C

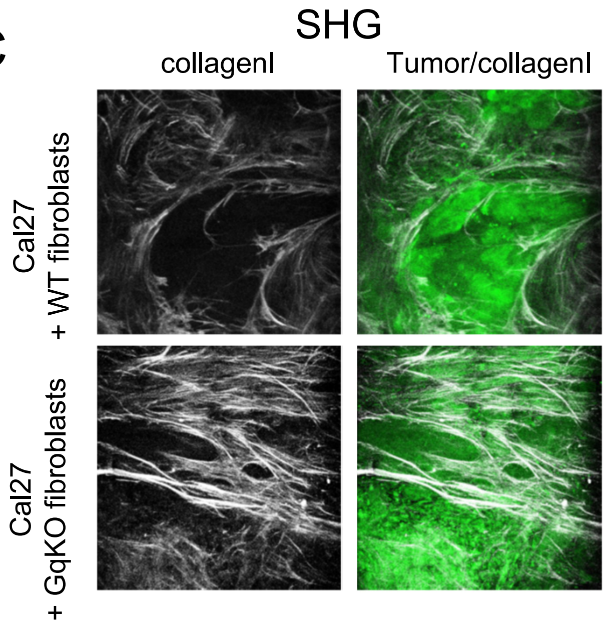

### Figure S7

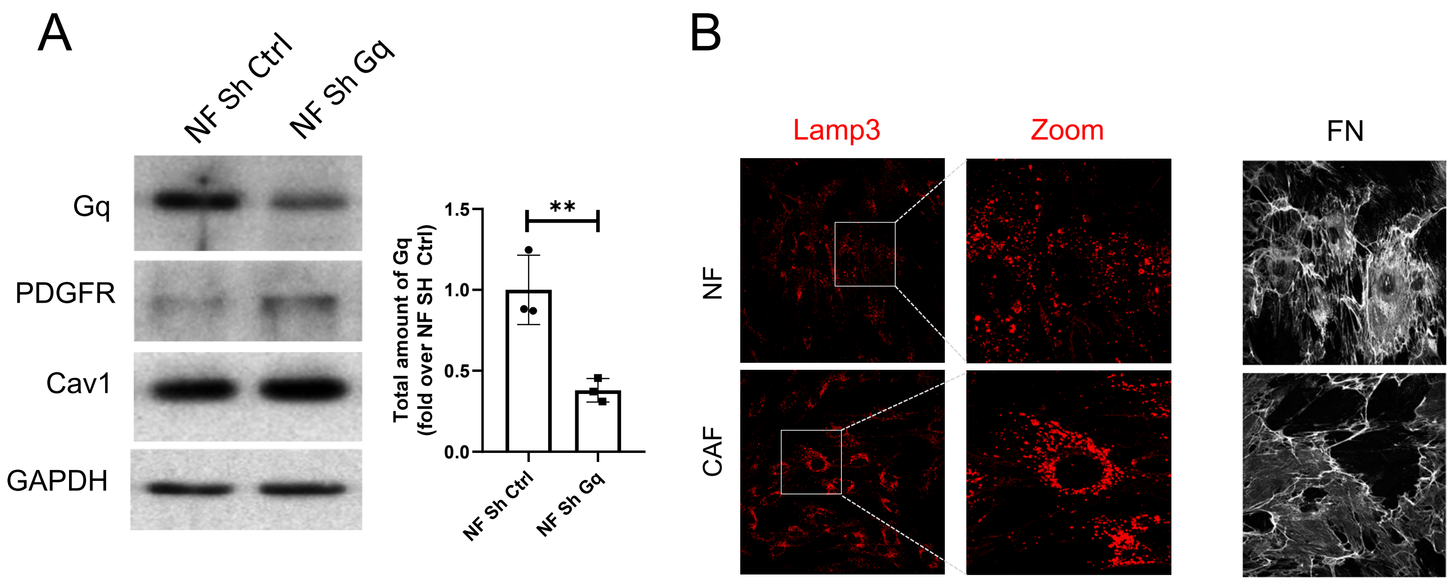
A
NF Sh Ctrl
NF Sh Gq
Gq
PDGFR
Cav1
GAPDH
Total amount of Gq (fold over NF SH Ctrl)
1.5
1.0
0.5
0.0
**
NF Sh Ctrl
NF Sh Gq
B
Lamp3
Zoom
FN
NF
CAF
